# Order invariance of fitness makes evolution predictable despite strong history-dependent effects

**DOI:** 10.64898/2026.09.29.755513

**Authors:** Shaili Mathur, Clare I. Abreu, Olivia M. Ghosh, Jonas Cremer, Dmitri A. Petrov

## Abstract

Environmental change is ubiquitous in nature, yet understanding its effect on the predictability of evolution remains a challenge. Previous work in microbes has shown that growth and fitness in a particular environment can depend strongly on the environments encountered previously. If such history-dependent effects are large, then any sequence of environments must be thought of as unique, and even a modest number of environments could generate a combinatorially large number of evolutionary paths. This would make evolution in changing environments difficult to predict. Here, we use cycles of three environments, which is the smallest number of environments which allow for comparisons of different environmental orderings – X →Y →Z →X vs X →Z →Y →X – to empirically test how environmental order affects the fitness of *S. cerevisiae* mutants. We measured the fitness of ∼900 adaptive mutants in distinct three-environment cycles, and find large history-dependent effects which cause fitness in individual environmental components to differ drastically across the two directions of the cycle. However, these effects tend to compensate for each other, resulting in similar overall fitness across the whole cycle. This is true even when we specifically conduct experimental evolution in both directions of the same three-environment cycle. Of the ∼450 mutants we isolated, only a small number of strains showed higher fitness in the direction in which they evolved. By splitting the contribution of each environment on fitness into its effect on the current and next component, we show that the invariance of overall fitness to environment order emerges because the effect of an environment on the next component is largely independent of the identity of the subsequent component, and can be captured by an additive model. Despite this general pattern of compensation, we do find instances of non-additivity, some of which are partly due to history-dependent effects lasting beyond two growth components. Overall, these results suggest that despite large history-dependent effects, fitness across a sequence of environments can be surprisingly predictable.

## 1 Introduction

One of the central goals of evolutionary biology is to understand the extent to which evolution can be predicted. There are abundant examples of convergent evolution in nature, suggesting that in any environment there exists only a limited set of available solutions, thus making evolution at least in part predictable. On the other hand, there are many reasons evolution could be unpredictable – evolution proceeds through inherently stochastic processes such as drift and mutation, which could be expected to hinder predictability of evolution, either by increasing the complexity of the environment and thus the number of possible solutions, or by introducing additional historical contingencies. A large body of experimental evolution studies has produced instances of consistent patterns of repeatability at the level of genes, pathways, and functional groups, supporting the view that evolution is predictable [1– 12], as well as cases of stochastic and unpredictable outcomes [8, 13–15]. Experimental evolution studies are particularly well suited to probe the predictability of evolution because they allow replicate populations to evolve under identical conditions. However, these studies have traditionally focused on evolution under relatively stable environmental conditions, often involving repeated growth on the same nutrient source. Microbes face environmental change in many settings, such as diurnal and seasonal environmental variations or repeated boom and bust cycles in nutrient availability, for example in a host gut or during a domestic fermentation process. Given that environmental variability is ubiquitous in nature, it remains unclear to what extent the observations from traditional experimental evolution can be applied to evolution in natural populations. Indeed, studies of evolution in fluctuating environments have found that fluctuations make evolution less repeatable [5, 16–18].

Furthermore, strongly fluctuating environments may be fundamentally different from more constant environments because organisms must adapt not only to multiple environmental states, but also to the transitions between them. Studies on microbial populations suggest that such transitions indeed pose a novel challenge for cellular growth separate from balancing multiple environments that constitute the fluctuation [19– 26], and fitness in fluctuating environments can deviate from predictions based on fitness measurements in constituent environments [16, 27, 28]. Moreover, on an evolutionary timescale, adaptations acquired in an earlier environment may influence which mutations are beneficial in subsequent environments due to epistatic or pleiotropic effects [14, 15, 29–31], causing populations exposed to different environmental orders to follow distinct evolutionary trajectories.

If environmental transitions can have large effects on fitness, we might expect that different orders in which the same environments appear could generate distinct evolutionary paths, even if the total exposure to the constituent environment components remains unchanged. For instance, consider the shortest cycle of distinct environments in which order can differ, X →Y →Z →X and X →Z →Y →X (Figure 1A). Large idiosyncratic fitness effects of environmental transitions could make these two sequences of environments very different conditions from each other.

**Figure 1:**
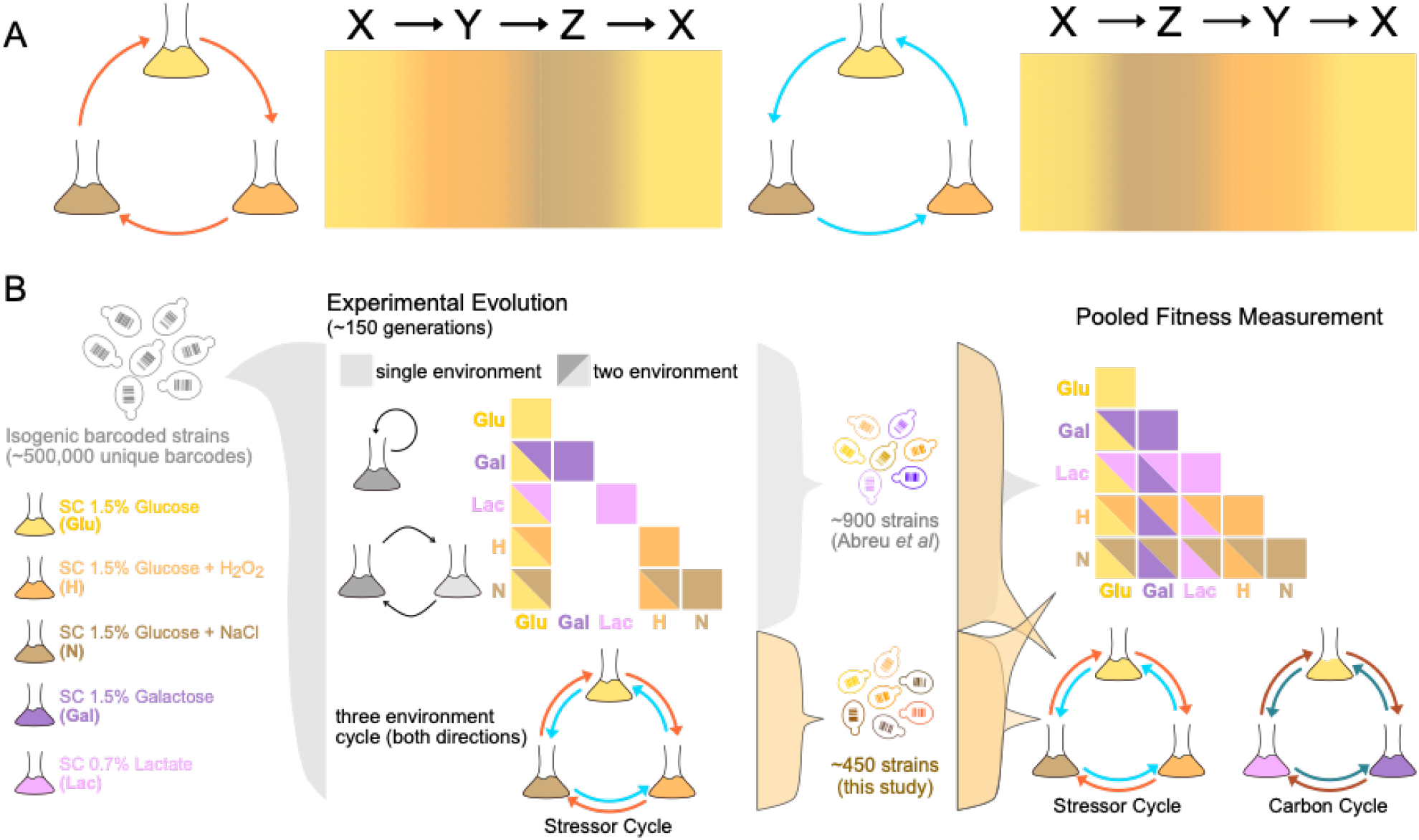
Schematic of Experimental Design. **(A)** Three-environment cycles have two different orderings (X→ Y →Z →X and X→ Z → Y→ X), which have different transitions, even if the mean time spent in each environment is the same. **(B)** We use a library of ∼900 experimental evolution mutants from one- and two-environment cycles, evolved from an ancestral population with ∼500,000 isogenic barcoded lineages. Strains from each condition were isolated, individually sequenced for barcode identity and pooled. ∼900 of these mutants were generated via evolution in one- and two-environment cycles and their fitness was measured in the five static environments and all possible two-environment cycles constructed from these environments in our previous study [27]. In this study an additional ∼450 mutants are generated via evolution in both directions of the three-environment cycle comprised of SC Glucose, SC Glu + NaCl, and SC Glu + Hydrogen Peroxide. We measured the fitness of these strains in three-environment cycles comprised of Glucose and the two stressors (NaCl and H_2_O_2_), as well as SC Glucose and the two alternative carbon sources (Galactose and Lactate). Additionally, we measured the fitness of strains generated from both directions of the three-environment cycles in all one, two, and three-environment cycles. Experiments from Abreu *et al*. [27] are shown in gray, measurements from this study are shown in orange.

If, however, history-dependent effects were primarily driven by only the current environment and the immediately preceding environment, this would simplify the three-environment cycles. Furthermore, if the effect of an environment (Y ) on future fitness were independent of the environment that follows it (X), rather than being an emergent effect arising from both of the environments in a fluctuation, then history-dependent effects might cancel out over time. To see this more clearly, we can decompose the fitness in environment X preceded by Y, s(X, Y→ X), into the effects of each environment and an interaction term,

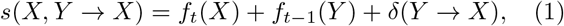

where f_*t*_(X) and f_*t−*1_(Y ) represent the fitness effects of environments independent of the other environment, and δ(Y → X) captures any interaction term. If there were only subtle history-dependent effects, then f_*t−*1_(Y ) and δ(Y→ X) would be small; if there were large but idiosyncratic history-dependent effects then δ(Y →X) would be large; if there were a constant effect of an environment on future fitness, independent of future environments, then δ(Y → X) would be small in comparison to f_*t−*1_(Y ).

In a three-environment cycle, each environment is the current environment and previous environment for one of the components. If δ(Y→ X) is small, then the overall fitness of a mutant in a three-environment cycle simplifies; for instance in environment X → Y → Z → X, overall fitness is 1/3*·Σ*_*E∈{X,Y,Z}*_ f_*t*_(E)+f_*t−*1_(E) ), which is invariant to the direction (order) of the cycle. As a result, overall fitness could become largely independent of environmental order. This would allow for a coarse-grained description of fitness in fluctuating environments over long timescales rendering evolution in fluctuating conditions predictable despite the history-dependent effects.

Here, we perform experiments in yeast, using a collection of DNA-barcoded *S. cerevisiae* strains to measure fitness in both directions of three-environment cycles (X →Y →Z→ X … and X→ Z →Y →X …), where cells are grown sequentially in batch culture on different media. We first measure the fitness of a collection of ∼900 high-fitness strains that were previously evolved in one- and two-environment cycles using the same media conditions used here [27], but never experienced the three-environment cycles we construct. Using this large collection of mutants allows us to quantify how much environmental order matters for fitness. We further analyze data from two-environment cycles to test the degree to which history-dependent effects are driven by idiosyncratic effects specific to an environmental pair.

Finally, we test whether environmental ordering affects adaptation by evolving strains in both directions of the three-environment stressor cycle. If strains fail to evolve direction-specific adaptations, this would suggest that environmental ordering has a limited influence on adaptation and that evolutionary outcomes may be predictable from simpler environments. Conversely, if strains gain adaptations specific to the direction in which they evolved, this would suggest novel adaptation to specific environmental orderings can occur rapidly, which may limit the predictive power of short-term fitness assays in fluctuating environments.

## 2 Results

To investigate how the order of encountered environments affects fitness, we first measured the fitness of a collection of ∼900 adaptive barcoded mutants that were previously generated through experimental evolution in constant or pairwise fluctuating environments [27]. Specifically, these mutants evolved in batch cultures containing one of five media components, i.e. synthetic complete (SC) glucose (Glu), SC galactose (Gal), SC lactate (Lac), SC glucose + NaCl (N), or SC glucose + hydrogen peroxide (H), or in cycles switching between pairs of these media conditions (Methods 4.1). Using this mutant collection, we probed fitness in two different three-environment cycles constructed from the components the strains were evolved in (Methods 4.3). The “carbon cycle” consisted of Glu, Gal, and Lac, while a “stressor cycle” consisted of Glu, N, and H (Figure 1B).

We measured fitness by pooling the barcoded mutant strains with the ancestor strain, and passaging this pool for six 48-hour growth cycles. The ancestor strain took up the majority (95%) of the initial culture so that competition is approximately between the ancestor and each mutant, rather than interactions between mutants. Cultures were transferred between environments via a 1:250 dilution (∼8 generations) every 48 hours, and the rest of the culture was sequenced to measure frequency at each time point. From frequency measurements of mutants and ancestor over time across batch growth cycles (Figure S1, S2), we model the fitness relative to the ancestor of a mutant i as

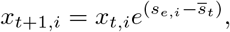

where x_*t,i*_ is the frequency of the strain at time t, s_*e,i*_ is the fitness of the strain in the environment between time points t and t + 1, and s_*t*_ is the integrated change in mean population fitness over the time interval. We define the fitness of the ancestral strain to be 0, and use the frequency change of the ancestor to infer s_*t*_ at time t. In multiple environment cycles, we jointly infer fitness in individual environment components (fitness during growth in one environment) and across the cycle (Methods 4.6).

### 2.1 Overall fitness in three-environment cycles is largely independent of ordering despite history-dependent effects

We compare fitness of the collection of ∼900 mutants from one- and two-environment evolution in the two directions of the three-environment cycles (Figure 2A,B). First focusing on the individual components (Figure 2 C-E, G-I), we find that fitness differs substantially across the two directions (mean absolute difference in fitness across the two directions of the cycle within the same component is 0.252), which is consistent with our previous findings that environmental history shapes fitness [27]. In contrast, when looking over the entire cycle instead of over individual components, we find that overall fitness in the two directions is generally well correlated (Figure 2F,J; mean absolute difference in fitness across the two directions of the cycles is 0.092).

**Figure 2:**
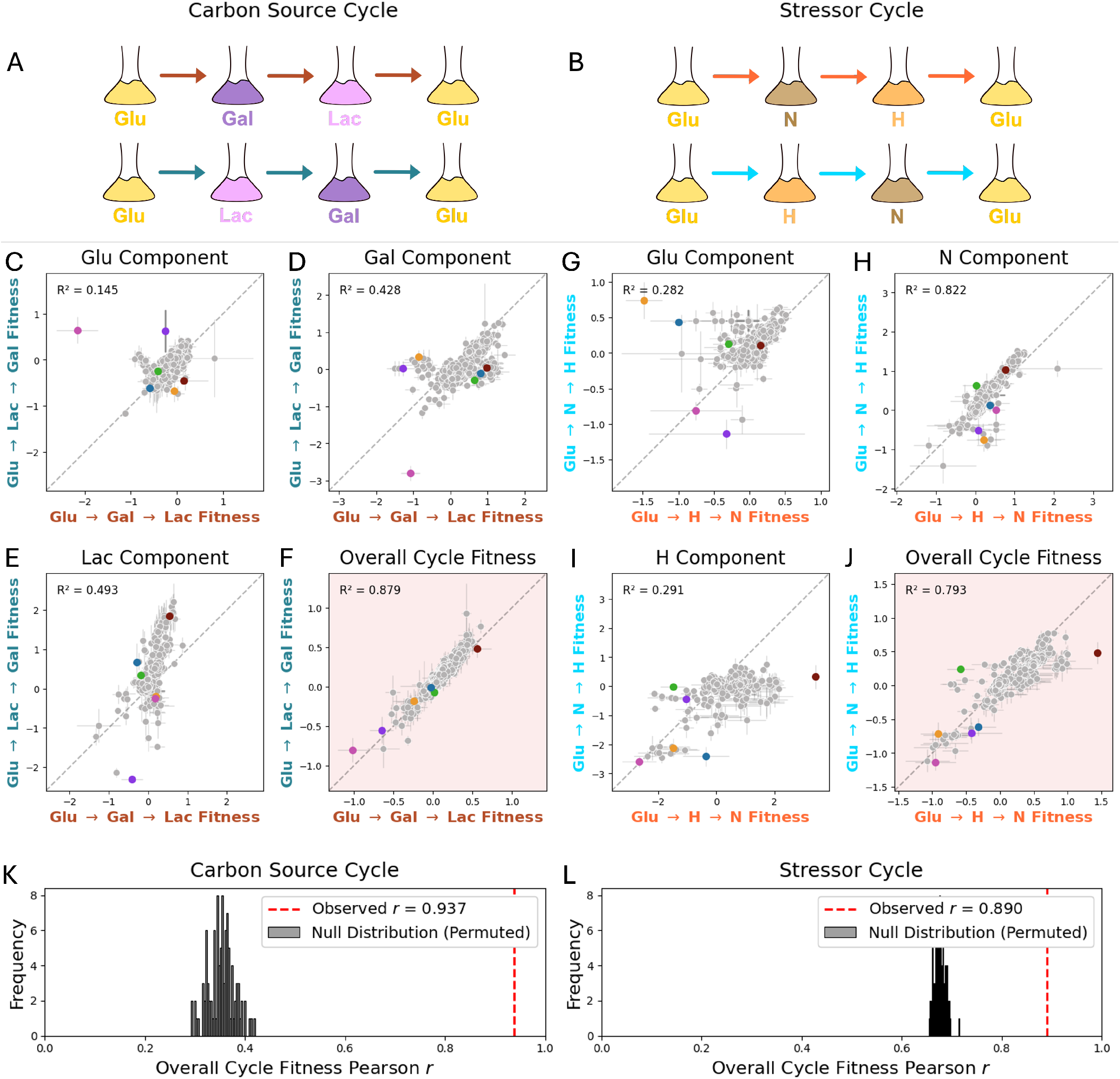
Fitness measurements in three-environment cycles of mutants evolved in one- and two-environment cycles. **(A)** Carbon cycle and **(B)** stressor cycles. **(C-E)** Fitness in each component of the three-environment carbon cycle compared across the two directions of the cycle. Strains differ in fitness across the two directions in components, indicating that there are large history-dependent effects. A selection of strains representing a range of behaviors are colored consistently across panels C-J. **(F)** Overall fitness in the two directions of the three-environment carbon cycle, calculated as the mean of fitness in the three environments. **(G-I)** Fitness in each component of the three-environment stressor cycle compared across the two directions of the cycle. Strains differ in fitness across the two directions in components, particularly in the glucose (panel **(G)**) and hydrogen peroxide (panel **I**) components, indicating that there are large history-dependent effects. **(J)** Overall fitness in the two directions of the three-environment stressor cycle, which is calculated as the mean of fitness in the three-environments. **(K**,**L)** Pearson correlation between overall fitness in the two directions of the carbon (panel **K**) and stressor (panel **L**) cycles are shown in red. The expected distribution of Pearson correlations between the two directions assuming that fitness between components is uncorrelated is shown in gray. This is calculated by permuting the fitness values across strains, maintaining the distribution of fitness values within each environment component.

We illustrate this by highlighting a few strains representing a range of behaviors in these environments. For instance, the fitness of the strain highlighted in pink in the Glu component of the carbon cycle is much higher in the Glu→ Lac →Gal direction than in the Glu→ Gal →Lac direction (Figure 2C) but this fitness difference is reversed in the Gal component (Figure 2D). Thus, the overall fitness averages to be very similar in both directions of the cycle (Figure 2F). Another example is the strain highlighted in orange, which has much higher fitness in the Glu →N→H direction of the stressor cycle in the Glu component (Figure 2G), but this advantage is compensated for in both the N and the H components (Figure 2H,I), in which the mutant is fitter in the Glu→ H→ N direction, again resulting in a very similar overall fitness. The maroon strain, on the other hand, is an exception to the general trend of similar overall fitness in the two directions of the fitness cycle. This strain has very similar fitness in both directions of the cycle in the Glu and N components (Figure 2G, H), but substantially higher fitness in the Glu→ H →N direction in the H component (Figure 2I), thus resulting in higher overall fitness in the Glu →H→ N direction of the stressor cycle (Figure 2J).

We use a permutation test to ask if the correlation in overall fitness is stronger than expected under the null hypothesis that the fitness of a mutant in each component of the three-environment cycle is uncorrelated with fitness in another component. We generate the null distribution by permuting the labels of strains within each environment, thus breaking correlations in fitness values between components but preserving the distribution of fitnesses for any given environment. We find a statistically robust effect in both cycles (Permutation test, p < 10^*−*3^) suggesting that over-performance in any given component in one direction of the cycle does generally trade-off with under-performance in another (or multiple other) components of the cycle (Figure S3I-J, S4). This pattern of compensation is present in both cycles with a larger effect in the carbon cycle than in the stressor cycle (Figure 2K-L). In a more detailed analysis across strains and conditions, we find that fitness behaviors in three-environment cycles depend on the conditions the strains evolved in (Figure S3). For instance, strains which evolved in the Glu/H and Lac in particular have large asymmetries in components. Strains which evolved in Glu/H are generally fitter in H when preceded by Glu (Glu →H →N direction) than when preceded by N (Glu →N →H direction), suggesting that they evolved specifically to the transition from glucose (no stressor) to hydrogen peroxide.

Taking these results together, we find that while fitness within a single component depends strongly on the previous environment, across the whole cycle fitness is often compensated such that overall fitness is invariant to the order of the environments. This invariance makes the overall fitness in fluctuating conditions more predictable as a result.

### 2.2 The contributions of each environment towards current and future fitness are largely additive

How can long-term fitness across the three-environment cycles be largely independent of direction, when there are substantial direction-dependent fitness effects in any given component environment? Considering fitness during an environmental switch, there are two distinct conceptual possibilities: (i) switching from one environment to another produces an entirely novel problem, and therefore fitness is an emergent property of the specific pair of environments, or (ii) there are additive effects of each environment on current fitness and fitness in the next environment component, such that the fitness contribution of each environment is independent of the other environment it is paired with.

More formally, we can decompose fitness of a mutant i measured in environment X preceded by Y, s_*i*_(X, Y → X), as

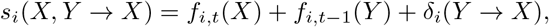

where f_*i,t*_(X) is the effect of environment X on the mutant’s fitness in the current environment, f_*i,t−*1_(Y ) is the effect of *previous* environment Y on the mutant’s fitness in the subsequent environment X that is independent of the specific identity of X, and δ_*i*_(Y→ X) is an emergent fitness effect due to an interaction between the two environments X and Y (Figure 3A). Importantly, f_*i,t*_(X) and f_*i,t−*1_(Y ) do not depend on the environment pair, and could also be used to model fitness effects in other switches involving these components, say Z →X or Y →Z. Under worldview (i) the expression for fitness is dominated by the interaction between X and Y, so δ_*i*_(Y→ X) would be large. Conversely, under worldview (ii), δ(Y→ X) would be small, so the effects of environments on fitness in the current and next environment do not depend on the specific environment pair in the fluctuation, and can instead be applied to any pair of environmental fluctuations.

**Figure 3:**
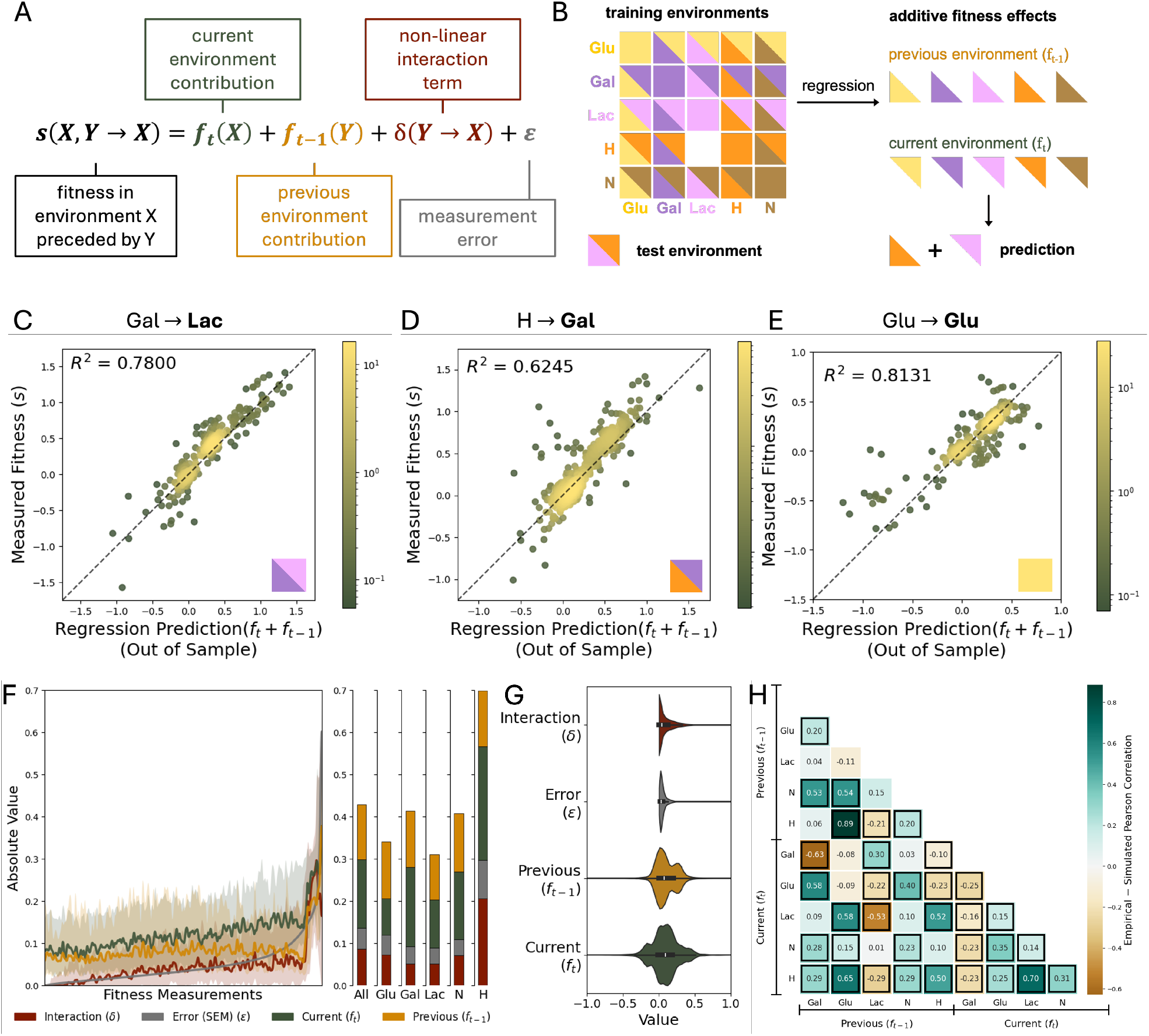
Regression separating generalized effects of environments on fitness in the current and next time point. **(A)** Model of generalized fitness effects of environments on the previous and current timepoint. For a given mutant, fitness in an environment X preceded by Y, s_i_(X, Y → X), is modeled as the effect of environment X on fitness when X is the current environment, f_*t*_(X), the effect of Y on the next environment, f_*t−*1_(Y ), an emergent interaction term, δ(Y→ X), and error (ε). Note that f_*t*_(X) and f_*t−*1_(Y ) are invariant to the other environment, i.e. the effects of X and Y independent of specific environmental transitions. **(B)** Schematic of ridge regression procedure. For each mutant, we jointly infer f_*t*_ and f_*t−*1_ for all environments, training the model on all data except the held out environment, which is then used to quantify prediction error. **(C-E)** Out of sample predicted fitness from the model fitting f_*t*_ and f_*t−*1_ as described in A-B compared to measured fitness. Points are colored by estimated density at that point (Gaussian KDE, Scott’s method), with lighter points indicating higher density. The panels show fitness measured in Lac preceded by Gal, Gal preceded by H, and Glu preceded by Glu (static glucose) respectively. **(F)** Absolute magnitude of contribution of the current environment f_*t*_ (green), previous environment f_*t−*1_ (orange), interaction term δ (red), and measurement error ε (grey) ordered by ε. The curves show the smoothed median and 0.25 and 0.75 quartiles, smoothed using Gaussian smoothing, with a rolling window of 100 data points with σ = 50. ε (measurement error) is estimated by the SEM of the fitness measurement. The SEM is subtracted from the prediction error to give an estimate for δ (clipped at 0). For cases in which prediction error is lower than SEM, the δ is clipped 0. Bars on the right show the mean values of f_*t*_, f_*t−*1_, δ, and ε across all the data, grouped by the current environment. **(G)** The distribution of f_*t*_, f_*t−*1_, δ, and ε across all the data as a violin plot. **(H)** Heatmap of Pearson correlations of f_*t*_ and f_*t−*1_ corrected for systematic bias from simulations (bias is *≈*+0.1 for current-previous environment pairs and *≈−*0.1 for previous-previous and current-current pairs). Coefficient pairs with p < 0.05 (FDR corrected, two-sided permutation test) and outside the 99% confidence interval from the simulated null are highlighted.

To empirically distinguish between these conceptual possibilities, we use fitness measurements of the same ∼900 *S. cerevisiae* mutants as above in the five one-environment cycles and nine two-environment cycles (Figure 1B). With multiple instances of any given environment being the current or previous environment, we are able to infer the generic fitness contributions of environments by fitting our data to the model s_*i*_(X, Y→ X) = f_*i,t*_(X) + f_*i,t−*1_(Y ) (Figure 3B). We construct this model separately for each mutant, by encoding the previous and current components of the fluctuating environment as binary variables, and jointly inferring the values of f_*t*_ and f_*t−*1_ across all environments using ridge regression. To test the performance of this model, we use leave-one-out cross validation (LOO-CV) across environments to generate a predicted fitness value for each fitness measurement (Figure 3B)(Methods 4.7). The prediction error from this model is due to a combination of interactions between the environments (δ) and noise in the data (ε). Thus, if this model is able to fit the data well, we can infer that δ is small.

We fit this model for all 23 environment switches using LOO-CV (Figure 3B), and found that overall the model improves fitness prediction substantially over a baseline prediction that does not take previous environment into account, indicating that there are generic environmental fitness effects on the previous and next environments which can enable prediction in previously unseen fluctuating environments. R^2^ is improved from 0.395 in the baseline prediction using the average fitness across switches with the same current environment, without taking the prior environment into account, to 0.621 over the entire dataset, with some variability across environmental switches (Figure S5-7). The performance of this model for three example environments that represent the range of model performance -Lac preceded by Gal, Gal preceded by H, and Glu preceded by Glu - is shown in Figures 3C-E. Generally predictions from this model are well correlated with the observed values, but there are also several cases with large differences between observed and predicted fitness.

To quantify the degree to which interactions between environments (δ) contribute to fitness, we analyze the errors from the out-of-sample predictions. Because this model fits only f_*t*_ and f_*t−*1_, prediction error is the sum of ε (measurement error) and δ. To estimate δ, we approximate ε by the standard error of the mean, i.e the expected magnitude of the measurement error in fitness and subtract it from prediction error (Methods 4.7). Simulations suggest that this is likely an overestimate of δ (Supplementary Materials Section 2, Figure S6). The magnitude of the inferred f_*t*_ (green), f_*t−*1_ (orange), δ (red), and ε (gray) for each data point (a single fitness measurement for a given mutant and environment component) ordered by ε are shown in Figure 3F as smoothed lines, with 50% inter-quantile ranges.

We find that f_*t*_ and f_*t−*1_ are similar in magnitude (green, orange in Figure 3F, mean magnitudes for f_*t*_ and f_*t−*1_ are 0.16 and 0.13), consistent with the large history-dependent effects we find comparing fitness in the same component across different directions of the cycle (Figure 2). In Figure 3F (right), the red columns can be interpreted as the expected value for δ, and the sum of the gray and red (δ + ε) as the upper bound for δ. Following this interpretation, we find that δ contributes 50% or less to the sum of the fitness effects for 77.6 - 93.0% of the data, and to less than 10% for 17.6-47.6% of the data. On average over the whole dataset, δ is estimated to be 16.96% of the fitness effects. Comparing the absolute values of δ, ε, f_*t*_ and f_*t−*1_ across the dataset (Figure 3F, G), we find that δ is generally a smaller value than f_*t*_ and f_*t−*1_, the inferred values for the generalizable additive fitness contributions. However, there are several cases where δ appears to have a substantial contribution, with some systematic variations across environments, such as larger values of δ in the H environment (Figure 3F).

With this model of fitness in environmental switches, we further investigated whether there are relationships across different parts of the growth cycle by analyzing the correlation structure in the fitness effects of different environments. Such trade-offs might be expected given previous work investigating relations between lag times, growth and survival in stationary phase, and could constrain fitness in dynamic environments. Consider the growth dynamics underpinning fitness in environment X preceded by Y : the fitness effect of the previous environment, f_*t−*1_(Y ), might be reflected in the early phase of growth in X rather than later growth phases, which might be better captured by the fitness effect of the current environment, f_*t*_(X). If proliferation in X (captured by f_*t*_(X)) traded-off with either survival in late stationary phase in the previous environment Y or with initial growth in X (captured by f_*t−*1_(Y )), then we would expect to see a negative correlation between f_*t*_(X) and f_*t−*1_(Y ). Conversely, if the same functional trait conferred fitness (dis)advantages across all parts of the growth cycle, then f_*t*_(X) and f_*t−*1_(Y ) would be positively correlated.

The estimated correlations between the f_*t*_ and f_*t−*1_ values are shown as a heatmap in Figure 3H, with negative correlations in brown and positive correlations in teal. We correct for a systematic bias (Pearson correlation of *≈*+0.1 in pairs of current/previous environment variables, and *≈−*0.1 in pairs of current/current and previous/previous environment variables; see Supplement Section 2, Figure S6) generated from this regression method. We find many instances of significant correlations between coefficients (highlighted in Figure 3H, Figure S6-7). For example, for the two stressors, the correlations between f_*t*_ and f_*t−*1_ are all positive, suggesting a cross-protective effect across the two stressors. However, looking across all environments there is a range of correlations, both positive and negative, for f_*t*_ and f_*t−*1_ values. Overall, we find no evidence of widespread trade-offs between different parts of the growth cycle, under the assumption that environmental effects on future fitness primarily reflect stationary and lag phases of the growth curve.

The totality of the data supports worldview (ii): the effects of environments are largely additive and generalizable. This is consistent with our observations across the three-environment cycles that while there are large short-term history-dependent fitness effects, the overall fitness across the whole cycle is invariant to the direction. However, note that additivity is not unbreakable and we do detect a number of cases in which the δ term is large, and there are substantial effects due to the idiosyncratic, environment-specific interactions between the current and preceding environments.

### 2.3 Compensatory memory constrains evolution in three-environment cycles

So far, our observations that overall fitness in three-environment cycles is largely order-invariant due to small fitness effects attributable to non-additive environmental interactions (δ) are based on mutants evolved in one- and two-environment cycles. However, we reasoned that if evolution were conducted in a specific three-environment cycle, adaptive mutants might exploit the specific order and non-additive interactions between environments. To test this hypothesis we carried out experimental evolution in the two directions of the three-environment stressor cycle (Glu →H →N and Glu →N →H) and asked if they adapt specifically to their “home” direction.

The founding populations of ∼500,000 isogenic bar-coded strains (Figure 1B) were evolved for 150 generations across 24 growth cycles in each of the two orderings and in two replicates, similar to the previous evolutions in one- and two-environment cycles [27]. We then isolated 110 mutants from each of these evolution conditions, for a total of 440 mutants, and measured their fitness in each of the three-environment cycles, as well as in 14 of the one- and two-environment cycles which can be constructed from the five environment components (15 cycles constructed from Glu, Gal, Lac, N and H, excluding Lac/H due to poor replicate-replicate measurements) (Figure 1B, Methods 4.2, 4.3).

First, we asked whether mutants evolved in the stressor cycle exhibit fitness behavior specific to their home direction. Looking at the individual fitness components and overall fitness (Figure 4A-D), we observe that most mutants isolated from either direction (blue, orange) do not appear to have adapted specifically to their three-environment direction, but instead cluster together similar to mutants evolved in one- and two-environment cycles (gray).

**Figure 4:**
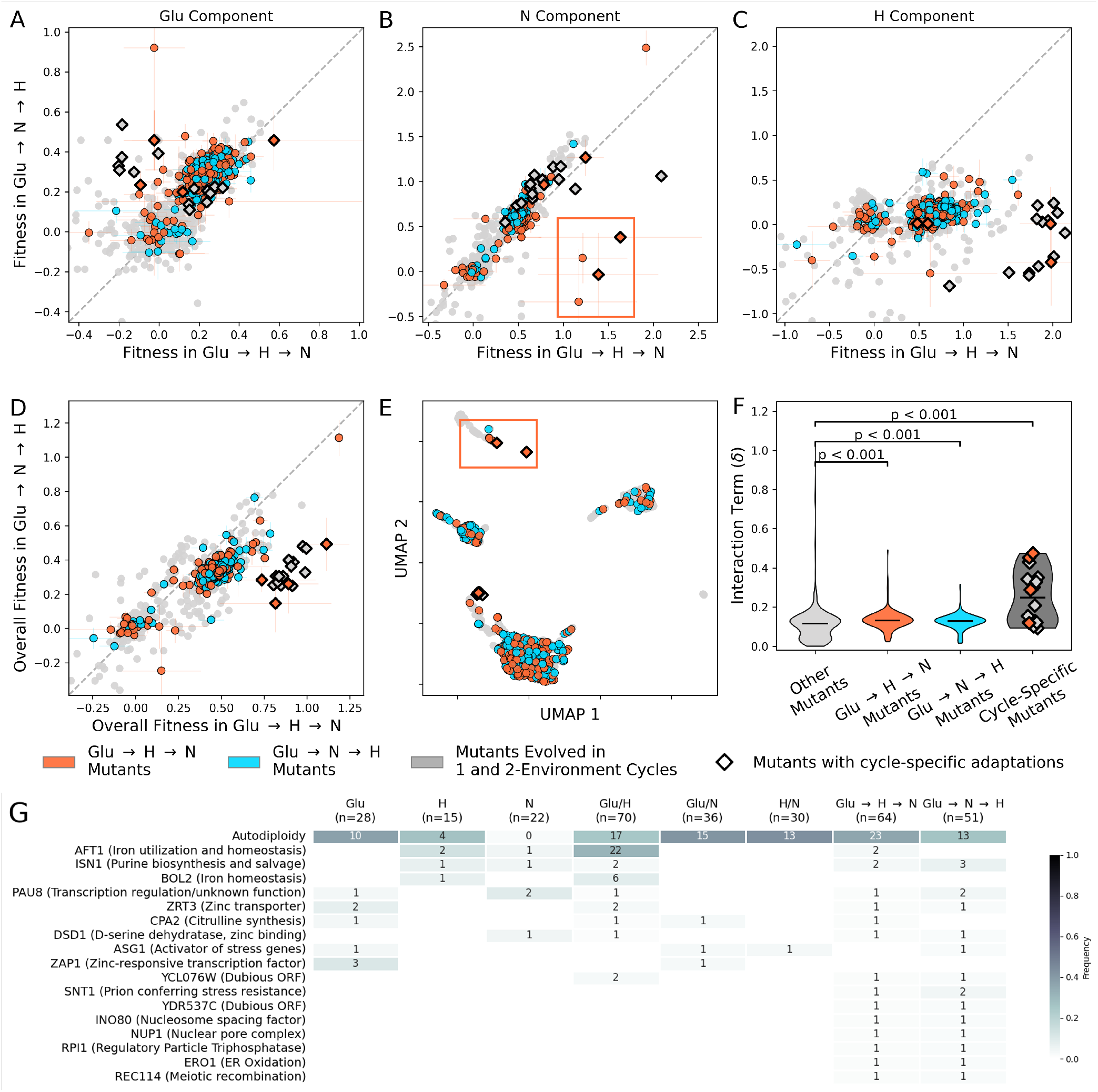
Evolution in three-environment stressor cycles. **(A-C)** Fitness of the evolved strains in the two directions of the three-environment cycles in individual components. Strains evolved in Glu →H →N are shown in orange, strains evolved in Glu →N →H are shown in blue, and strains evolved in one- and two-environments are shown in gray for comparison. Error bars for the three-environment cycle mutants show standard error of the mean (SEM) across three replicates. Strains showing higher overall fitness (fitness difference > 0.45 and overall fitness in Glu →H →N > 0.7) in the Glu →H →N cycle are highlighted with diamond markers; these strains are from Glu→ H→ N (n = 4), Glu/H (n = 10), Static H (n = 3) or Glu/Lac (n = 1). Strains from Glu →H →N showing specific adaptations to the H →N transitions are highlighted in the orange box in B and E. **(D)** Overall fitness in the cycle, calculated as mean fitness in the three components. **(E)** UMAP visualization of the fitness of all strains across all environments excluding the three-environment stressor cycle. Colors and markers are consistent across panels B-E. **(F)** Mean interaction terms δ for mutants across the components of the stressor cycle (Glu, H, N), grouped by their evolutionary origin. Highlighted strains with high fitness differences are shown in dark gray. p-values are obtained using a permutation test with n = 10, 000, and suggest that mutants in the Glu →H →N cycle have higher δ terms. **(G)** Most common genomic changes in stressor condition mutants from whole-genome sequencing, including autodiploidy and gene targets. Mutants from three-environment conditions share mutations in the most common gene targets with strains from all other conditions.

However, four mutants from Glu →H →N (orange mutants highlighted with diamond markers in all panels in Figure 4) gained fitness specific to their evolution direction. Two of these strains appear to have gained this asymmetric fitness advantage through adaptation specifically to high salt in the N component following H_2_O_2_ exposure in the H component(Figure 4 B). Interestingly, mutants from the two-environment H/N cycle have not evolved this ability, despite having the same H to N transition in their home environment. Notably, the outlier mutants with Glu →H →N specific adaptation are among the highest fitness strains in this cycle. This suggests that while rare (4/220 strains from Glu →H →N showed high fitness specific to the cycle direction), evolution specific to the cycle direction is one of the more effective adaptive strategies.

Additionally, we observe that most strains from both directions of the evolution are fitter in the Glu→ H →N direction than the Glu→ N→ H direction, particularly in the H component (Figure 4C). This suggests that most adaptations are more beneficial when transitioning from Glu (lower stress) to H (oxidative stress) than when transitioning from N (osmotic stress) to H, regardless of the cycle direction in which they were selected. Nevertheless, many of these strains appear to have lower fitness in the Glu component in the Glu →H→ N direction (Figure 4A), thus resulting in only a modest fitness difference in the overall cycle (Figure 4D). Using a permutation test, we find that the over-all fitness of these mutants in the three-environment cycles is more similar than in the components (Figure S8), similar to mutants from other evolution conditions (section 2.1).

Next, we examined how strains evolved in the three-environment cycles performed across all other environments. We visualize the behavior of strains across the collection using UMAP (Figure 4E), and find that strains evolved in the three-environment cycles are generally well distributed across the entire space. Reasoning that similar patterns of fitness across several environments indicate a similar underlying phenotype [32], this suggests that these mutants generally explore a similar phenotypic space as the mutants from one- and two-environment cycles. However, two of the four strains which show cycle-specific adaptations to Glu→ H→ N do so by adapting specifically to N in the Glu → H →N cycle, which no other strains appear to have done. These two strains are clustered with each other away from other strains (box, Figure 4 B, E), suggesting that these two strains have similar phenotypes to each other based on their overall behavior in all other environments.

In order to investigate the genomic basis of evolution in the three-environment cycles, we sequenced whole genomes of 115 strains from the three-environment cycles, and also tested strains for auto-diploidization (Figure 4G). We observe a genomic signature of environment-specific adaptation in the three-environment cycles. For instance, strains with SNPs in genes that are hit multiple times in the three-environment cycles but never in other cycles have higher fitness in three-environment cycles than other strains (Figure S9). There is no clear mechanism based on known functions of these genes (multi-hit genes in both cycles shown in Figure 4G, multi-hit genes from individual cycles listed in Table S1). This set of strains includes two of the outlier strains with adaptations specific to the Glu → H → N cycles.

We also find that there are two occurrences of mutations in *AFT1*, which is part of an iron-sensing pathway involved in oxidative stress response. This mutation occurred only in Glu → H →N mutants, not in Glu →N →H mutants, and was a common mutation in mutants from the two-environment Glu/H cycle, but did not occur in mutants from the H/N cycle. Interestingly, one of the two Glu →H →N strains with an *AFT1* mutation shows a much higher fitness in Glu →H →N than in Glu →N →H. This suggests that these are adaptations specific to transitioning to oxidative stress from a lower stress environment, but not from osmotic stress.

Looking over the whole collection of mutants from three-environment cycles however, we find that evolution remains largely constrained to generate fitness that is similar in the two directions of the cycle. This is in line with the order-invariant fitness and the low environmental interaction term (δ) we inferred in mutants from one- and two-environment cycles. Applying this analysis to the three-environment mutants, we find that most three-environment mutants also have small δ values. As expected, the outlier strains in Glu →H →N that have specifically adapted to their home three-cycle direction do have high δ values (Figure 4F). Given that these strains were among the fittest in this environment, it is possible that in a longer evolution they would have taken over the population, thus breaking the cycle direction symmetry constraint in longer term.

### 2.4 The length of environmental memory can exceed one growth cycle

One possible mechanism for a large interaction term (δ in Section 2.2) is long-term environmental memory. If memory exceeds beyond the immediately preceding growth cycle then the terms f_*t*_ and f_*t−*1_ of the current and previous environment cannot capture the effects of a third environment. Note that there are eight generations of growth in each growth cycle and hence over 99% of the transcripts, proteins and other cellular material should be diluted out of the cell by growth over each 48-hour growth cycle. This suggests that *a priori* such long memory is unlikely.

Nonetheless, we probed long term environmental memory by comparing the two-environment cycle to three-environment cycle fitness (Figure 5A). A difference in fitness measured in the last environment component, X, in Z →Y →X (three-environment) and in X →Y →X (two-environment), is a measure of the degree to which the first environment affects fitness (Figure 5A). We term this “T-2 memory”. Correspondingly, fitness differences attributed to a different environment immediately before the current environment, such as fitness differences within components across the directions of three-environment cycles, is termed “T-1 memory”.

**Figure 5:**
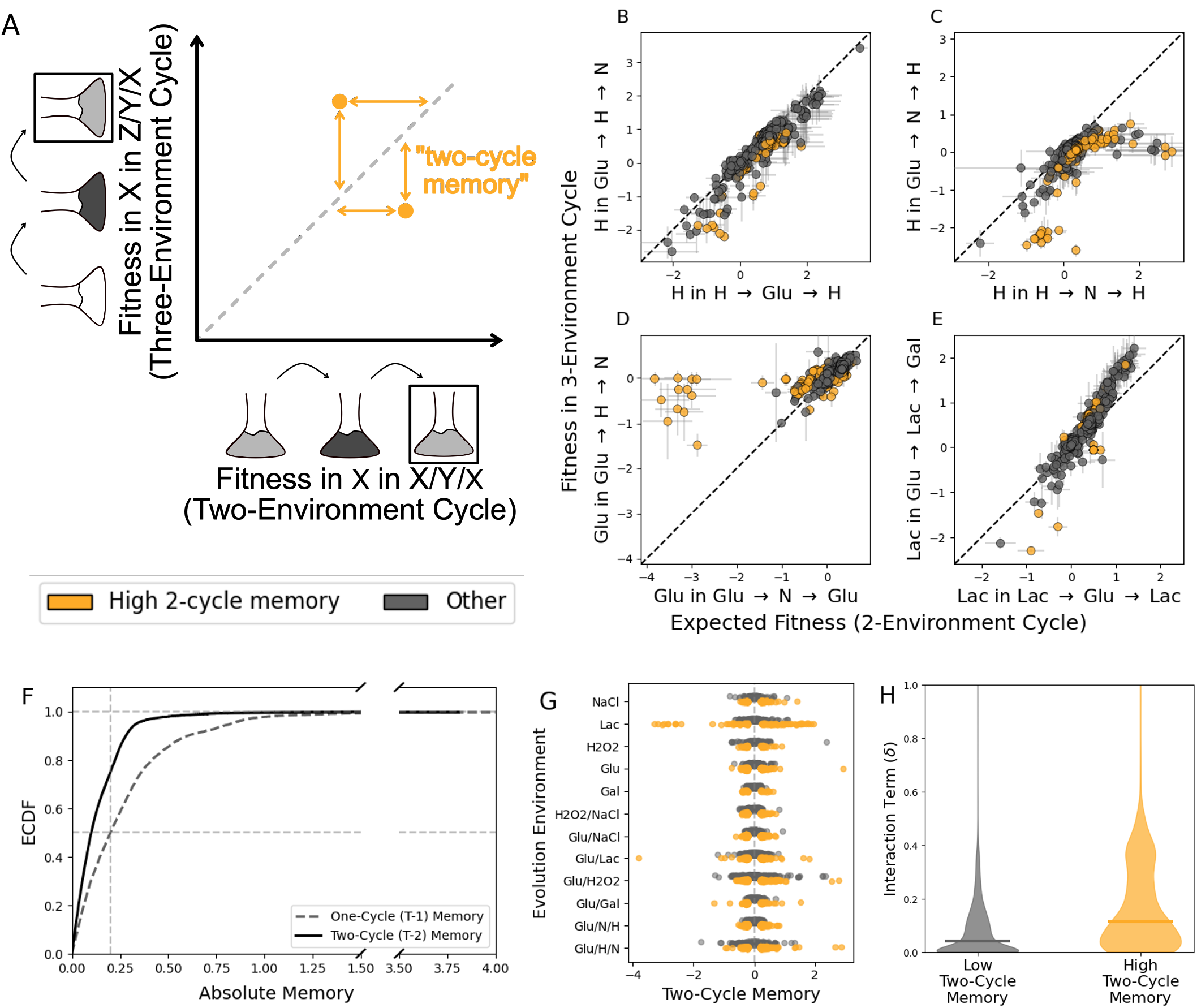
Evidence of longer term history-dependent fitness effects by comparing two and three-environment cycle fitness. **(A)** Schematic representation of plots showing longer-term memory effects. Comparing fitness in a three-environment cycle (i.e. fitness in X component in Z →Y →X) with fitness in the two-environment cycle (i.e. fitness in X component in X →Y →X) with the same previous environment isolates the effect of an environment on fitness two growth cycles in the future. We term the fitness difference between three-environment fitness and two-environment fitness “two-cycle (T-2) memory”. Measurements with substantial T-2 memory (T-2 memory > 0.2, p-value *≤* 0.138, in the smallest 31% of p-values) are highlighted in orange (p-values calculated using Welch’s t-test and Storey’s π_0_; see Methods 4.8 for further details). **(B-E)** Fitness in each three-environment component is compared to fitness in the two-environment component which has the same preceding (and current) environment. Each point represents the mean fitness of a mutant in the given environment, error bars are given by the standard error of the mean (SEM). We show a subset of environment components with a range of different behaviors: **(B)** H in Glu→ H →N, with low T-2 memory, and **(C)** H in Glu→ N→ H, **(D)** Glu in Glu →H →N, and **(E)** Lac in Glu →Lac →Gal all with several mutants showing substantial T-2 memory. **(F)** ECDF showing absolute value of T-2 memory and T-1 memory. T-2 memory (black, solid), while substantial, is generally smaller in magnitude than one-cycle memory (grey, dashed). **(G)** Two cycle memory is structured by the evolution condition strains were evolved in. In particular, strains from Glu/H, Glu →H→ N, and Lac appear to show large T-2 memory. **(H)** Strains with higher T-2 memory have larger interaction terms (δ), suggesting that longer-term memory may contribute to non-additive effects.

For each fitness measurement in three-environment cycles (n = 14, 640), we compare fitness measured in three-environment cycles to the corresponding two-environment cycle (Figure 5B-E, Figure S10). We estimate that ∼ 31% of these comparisons are true positives (Welch’s t-test, Storey’s π_0_ = 0.69), and colored cases of substantial T-2 memory (T-2 memory > 0.2, p-value *≤*0.138, bottom 31% of p-values; n = 2774, 18.95% of measurements) in orange. For many environments, such as H in N →Glu →H (Figure 5B), fitness in two and three-environment cycles is similar. However, there are several environments in which there are striking deviations from the expected fitness from two-environment measurements. For instance in H in Glu →N →H, mutants evolved in the Glu/H fluctuating cycle, have much lower fitness than expected (Figure 5C), and several mutants evolved in Lac have much higher fitness in Glu in Glu →H →N than expected (Figure 5D). Interestingly, high fitness in the Lac component in the Glu/Lac two-environment cycle is correlated with high T-2 memory in Lac in Glu → Lac → Gal (Figure 5E). See Supplement Figure S10 for corresponding plots in all environments.

Across all mutants and environments, the T-2 memory is smaller in magnitude than T-1 memory (Figure 5F); 24.9% of measurements exceed T-2 memory of 0.2 (corresponding to ∼20% frequency change per cycle relative to ancestor), in comparison to 50.1% of the data points with T-1 memory greater than 0.2. Interestingly, these effects are structured by the evolutionary origin of the strains; strains evolved in Glu/H, G→ H →N, and Lac show the largest two-cycle memory effects (Figure 5G).

Grouping measurements by the magnitude and significance of long-term memory, we observe that cases with higher longer-term memory effects disproportionately have higher interaction terms (δ) from the model in 2.2 (Figure 5H, Figure S10). Overall, our measurements of a large number of mutants in three-environment cycles show that there is widespread onecycle memory from one growth cycle to the next and a secondary but occasionally substantial effect of two-cycle memory which contribute to larger interaction terms between environments.

## 3 Discussion

To investigate the predictability of evolution in fluctuating conditions, we studied how the order of environments affects evolutionary outcomes. If each ordering of environment components created functionally different regimes, then each sequence of even a small number of environments would result in a combinatorially large number of unique environmental paths, which would make it very difficult to predict evolution in fluctuating environments. We considered three-environment cycles, which are the shortest cycles in which environments can be ordered differently. The two possible directions of a three-environment cycle (X →Y →Z →X and X →Z →Y →X) have the same mean time spent in each environment, but different transitions from one environment to another. We constructed these environment cycles in a batch culture setup, re-diluting cultures into changing media, with the growth dynamics resembling boom-and-bust cycles in natural environments.

We first measured the fitness of ∼900 strains generated via experimental evolution in several one- and two-environment cycles. We assessed their fitness in two three-environment cycles: a “carbon” cycle where cells are exposed to different carbon sources and a “stressor” cycle where cells are exposed to varying stress levels. We found that there were large differences in fitness across the two directions within individual components of the cycle, i.e. history-dependent fitness effects. Strikingly however, these history-dependent effects largely cancel out such that overall fitness across the two orderings is similar. We further tested the robustness of this result by evolving strains in the two orderings of the three-environment stressor cycle and measuring the fitness of ∼450 isolates in both directions of the cycle. We found that strains largely did not evolve order-specific adaptations, with some notable exceptions in the Glu → H → N direction.

We reconcile the invariance of overall fitness to the order of environments with large history-dependent effects within components by reasoning that fitness in a fluctuating environment can be decomposed into additive effects. In this case, fluctuating fitness s(X, Y→X) can be written as the sum of the fitness effects of the current environment (f_*t*_(X)) and the previous environment (f_*t−*1_(Y )). Any interaction term which would capture an emergent effect specific to the pair of environments (δ(Y → X) = s(X, Y → X) *−* f_*t*_(X) *−* f_*t−*1_(Y )) would be small on average. Importantly in this model, the previous and current environment effects are general to all environments, and could be used to predict fitness in entirely different environment pairs, e.g. X→ Z or Z →Y. Using two-environment fitness data, we test predictions of a model that only captures additive effects (f_*t*_, f_*t−*1_) of environments, and find that fitness in held-out environments is well predicted, suggesting that interaction terms capturing the emergent effects on fitness are on average small.

Taken together, these results present a surprisingly simple framework for predicting fitness in fluctuating environments, even amid large history-dependent fitness effects. Of course, natural settings and populations are likely much more complex than what we can recreate in the lab, with more diverged strains or species, a larger degree of community interactions, or more complex or stochastic environmental fluctuations. Further studies are therefore crucial to understand the generality of these effects in a broader range of environments and strains. Nevertheless, the predictability of environmental memory across a large number of strains and environments is striking.

From a physiological perspective, that fitness in a fluctuating condition can approximately be decomposed into the additive effects of current- and previous-environment contributions is surprising. In our batch culture experiments, cultures repeatedly grow, reach saturation, and are then diluted into fresh media. Upon dilution into fresh media, cells typically enter lag phase, a period of little to no growth which has been shown to contribute substantially to fitness variation [33]. Recent physiological studies have further established a complex dependence of lag phase on the cellular regulation of proteome composition and metabolic fluxes [23, 34–37]. This suggests that lag times would be idiosyncratic, depending on the specific strains and environments. Memory effects could arise because the physiological state of a strain at the beginning of a growth cycle depends on the environment it was previously in, and therefore the extent to which it must rewire itself to resume growth should depend on environmental history. Despite the complexity of these processes, however, our results imply that the fitness effects of cellular rewiring required for growth are approximately an additive function of the pre- and post-switch environments. The generality of the effects we observe across fluctuating environments also suggests that these changes occur in a fundamentally low-dimensional space, effectively driven by a small number of traits. Although this is not apparent given the complexity of cellular growth, such an intuition is consistent with several previous studies which have shown that genotype-phenotype-fitness maps can often be captured by low-dimensional models [32, 38].

We can use the correlational patterns in the inferred additive fitness effects of each environment to elucidate the extent to which there are trade-offs or synergies between different stages of growth [39–41] in our system. Some previous studies suggest that growth and lag-phase can trade-off with one another [35, 42]. Other work shows that lag-times can be tuned by allocation towards unneeded proteins, and can thus be shortened without sacrificing growth rates [34]. In our model, we consider the primary effects of the previous environment (f_*t−*1_) to be on lag phase and the current environment (f_*t*_) to be on exponential growth. When analyzing the correlations across mutants between the fitness contributions of a given environment on the next cycle (f_*t−*1_) and the current environment (f_*t*_) we see a range of correlations. In stressor environments, we see a pattern consistent with cross-protective effects [43– 46], possibly due to global stress response: strains that have an advantage in early growth coming from a stressor environment maintain this advantage during growth in a stressful environment. For carbon sources, we do not observe a uniform pattern — there are some cases where there appears to be a strong trade-off between early and exponential growth, as well as other cases where mutants with an early growth advantage maintain their advantage through the exponential growth phase. Our analysis hints at possible relationships between different growth stages measured at scale, and measuring effects on lag times or growth directly could be an avenue for future studies.

The invariance of fitness to the order of the environments suggests that adaptations specific to the environmental transitions, which could be interpreted as anticipatory behavior, are difficult to evolve. However, putatively anticipatory behavior has been engineered in *E. coli* [47] and several examples have been discussed in the context of natural settings, including predictable daily changes in light and temperature, seasonal changes, and the order in which nutrients or stressors occur [20, 21, 24, 25, 48]. Three-environment cycles are temporally structured and therefore predictable, making this setting ripe for the evolution of novel anticipatory behavior. From our experimental evolution experiment we found there are a limited number of instances where mutants from the Glu →H →N cycle showed significantly higher fitness effects specific to their home direction, suggesting that evolution of novel anticipatory behavior is possible even in the short term. Additionally, most strains have slightly higher fitness in the Glu →H →N than in the Glu →N →H cycle. Previous studies [25] suggest that osmotic stress often precedes oxidative stress in fermentation environments, so the Glu →N →H cycle might be considered more akin to a natural environment for *S. cerevisiae*. Therefore, easily accessible mutations could be biased towards increasing fitness in the Glu →H →N cycle, which is more different from the “natural” environment due to a global epistasis-like effect [49–51].

The environmental memory in two and three-environment cycles we observe also raises questions about the length, mechanistic origins, and universality of history-dependent effects. By comparing data from three-environment measurements to two-environments measurements, we asked whether memory effects can last longer than one growth cycle, or 8 generations in this experimental design. We find that memory effects are diminished after one cycle but appear to be non-negligible, and we estimate that 31% of our measurements reflect nonzero effects. We assume that history-dependent effects arise largely when cells are moved directly into the next environment in the cellular state they finished at in the previous environment, which is highly environment-specific. This picture cannot explain longer memory effects however, as virtually all proteins from two growth cycles before the current one would be diluted over the course of growth. These longer-lasting memory effects suggest that other mechanisms may be at work. These could include physiological processes like epigenetic changes or prion-based memory [52–56], or broader eco-evolutionary phenomena such as frequency-dependent fitness effects [57–60]. Further experiments would be needed to confirm this. Connecting cell physiology to growth and fitness in fluctuating environments is a fundamental challenge in microbial evolution. We used experimental evolution and high-throughput fitness measurements in differently ordered environments to show that fitness in fluctuating environments is highly predictable despite pronounced history-dependent effects. This predictability is underpinned by decomposing fitness additively into current and previous environmental contributions. Exploring the physiological mechanisms that lead to this simple fitness decomposition and the extent to which these results can generalize to complex natural settings provides an avenue to better understand how organisms adapt to a dynamic world.

## 4 Materials and Methods

### 4.1 Collection of Mutants from One- and Two-Environment Cycle Evolutions

We used a collection of 889 barcoded mutants from Abreu et al. [27] which were evolved in 5 one-environment cycles: SC Glucose (Glu), SC Glucose + Hydrogen Peroxide (H), SC Glucose + NaCl (N), SC Galactose (Gal) and SC Lactate (Lac), and 5 two-environment cycles: Glu/Gal, Glu/Lac, Glu/H, Glu/N, and H/N. Barcoded pools were sequenced over the course of the evolution, and isolated strains were sequenced to estimate their fitness in the evolution environment. Pooled mutants were high-fitness strains from their home evolution environment.

### 4.2 Evolution and Strain Isolation in Three-environment Cycles

We evolved strains in two three-environment cycles: Glu →H →N and Glu →N →H. We used the same protocol to evolve strains, isolate strains and measure fitness as in [27], and briefly describe it here. The founding population was previously created by integrating a genomic barcode into BY4709 [61]. We confirmed the presence of ∼500,000 unique barcodes before beginning the evolution experiment by sequencing the barcode region.

SC 1.5% Glucose base was made using 2g Drop-out Mix Complete (US Biological) and 6.7g yeast nitrogen base (RPI) and 15g glucose per liter. The stressor conditions N and H respectively were made by supplementing to 0.5 M NaCl (RPI), or hydrogen peroxide to 2.5mM H_2_O_2_(Ward’s). All media was prepared before the evolution experiments and stored at 4^*°*^C, and warmed to 30^*°*^C the day before use. We found that H_2_O_2_ decreased while the media was stored to approximately 60% of the original concentration using the MQuant peroxide test kit.

Evolution experiments were conducted in batch culture; cultures were grown in 100 mL of media in 500 mL DeLong flasks at 30^*°*^C, shaking at 223rpm, for 48-hours. At the end of each growth cycle 400 µL (∼5 *×*10^7^ cells) of the culture was diluted into fresh media, and 1 mL of the culture mixed in equal quantities with 50% glycerol, and stored at -80^*°*^C. We stored the remainder of the culture for barcode-sequencing by centrifuging the culture, resuspending the pellet in sorbitol solution (0.9M sorbitol, 0.1M Tris-HCl (pH 7.5), 0.1M EDTA (pH 8.0)), and freezing at -20^*°*^C. We evolved strains for 24 total growth cycles (48 days).

We sequenced the barcode region at several time points over the course of the evolution (see 4.4), and used FitMut2 [62] to infer the fitness of each lineage. To isolate strains, we chose an intermediate timepoint at which there were apparently adaptive mutants which had risen in frequency, but no mutants had reached a very high frequency, thus allowing us to isolate a number of adaptive mutants. 50 µL of the frozen stock (∼ 10^6^ cells) was diluted into 500 µL phosphate buffer solution (PBS) plus 1 µL of propidium iodide to stain dead cells. Live cells were sorted into 100 µL of YPD in 96-well plates using a flow cytometer. We sorted into five plates for each of the four evolution replicates.

Sorted cells were grown at 30^*°*^C without shaking for 3 days. The saturated culture was inoculated into fresh YPD media and used for sequencing to identify the bar-coded strains. The remainder of the culture was stored in 25% glycerol and stored at -80^*°*^C. We then sequenced each isolated strain to identify barcodes.

We selected adaptive barcodes for further study from isolated strains if they met any of the following criteria: (1) inferred fitness was 0.51 or greater; (2) the inferred probability of being adaptive was greater than 0.5, or (3) strains reached a final frequency of 10^*−*5^, which is approximately five times greater than their initial frequency. We eliminated barcodes which met these criteria for multiple conditions, including the one- and two-cycle evolutions, as these were likely preexisting adaptive mutations in the founding pool. We also included an additional set of weakly adaptive strains (with inferred fitness between 0.125 and 0.5) for each replicate. The final pool had 440 mutants: 220 mutants from Glu →N →H and 220 mutants from Glu →H →N. We pooled saturated cultures of each strain in approximately equal volumes for pooled competition assays. See Abreu et al. [27] for further details on these methods.

### 4.3 Fitness measurements in three-environment cycles

We measured the fitness of the strains from one- and two-environment cycles generated from Abreu et al. and the strains we evolved in three-environment cycles in a single competitive pool. We measured fitness in three replicates in four three-environment cycles: Glu →H →N, Glu →N →H, Glu →Gal →Lac, and Glu →Lac →Gal. The environments for fitness measurement were the same as those in which we did evolution experiments; pools were grown in 100 mL cultures in 500 mL flasks at 30^*°*^C, shaking at 223 rpm.

We followed the same protocol as in [27] to measure the fitness of each of the∼ 1300 strains in the pool. Briefly, the pooled culture consisted of the ancestor strain with a digestible barcode, the barcoded mutants, barcoded “neutral” strains which have no functional mutations from the ancestor, with the ancestor and neutral strains taking up most of the initial culture. Thus, the environment is largely expected to be set by the ancestor strain, and most pair-wise competition for any mutant strain can be assumed to be with the ancestor. We grew overnight pre-cultures in SC 1.5% Glucose of the ancestor strain, barcoded pool and barcoded “neutral” strains separately. We mixed these cultures weighted by OD600 at a 93:5:2 ratio. These cultures were passaged through each sequence of environments, diluting 400 µL of the culture into fresh media at the end of each 48-hour growth cycle, as in the evolution conditions. At each transfer, the remainder of the culture was frozen for barcode sequencing. We then sequenced each time point to measure the frequency of each barcode (Section 4.4).

These fitness measurements were performed concurrently with measurements in Abreu et al., minimizing batch effects from variation in media, growth conditions, pre-cultures, or other similar factors.

### 4.4 Barcode amplicon sequencing

We followed the same protocol as in [27] to sequence the barcode region of each sample. We extracted genomic DNA from each sample using the Norgen Fungi/Yeast Genomic DNA Isolation 96-Well Kit (#27350), using about 10^9^ cells per sample (10% of our original cultures), and measured DNA yield using a Biotek fluorescent plate reader averaging 4 µg per sample.

We amplified barcoded in two steps, checking reactions on a gel for∼ 50 randomly chosen reactions at each step. We used the same first-step primers described in Kinsler *et al*. [32]. The first-step primers had short inline indices which were used to mark samples, which we use in combination with second-step primers to multiplex the samples on the sequencing lane. We then diluted the first step PCR product 500-fold for the second-step reaction. The second step PCR used Illumina UD Indexes (from IDT), with unique pairs of forward and reverse primers for each sample. We performed PCR-reactions using Q5 master mix in 50 µL reactions. After purifying samples and measuring final DNA concentration (∼100 – 200 ng µL^*−*1^), we sent samples to Admera Health for sequencing, using 2*×*150 paired-end sequencing using HiSeqX or NovaSeqX. We used this protocol for sequencing both evolution timepoints and short-term fitness assays. For fitness assays, we used ApaLI to digest the barcode of the ancestor strain, so that most of our sequencing reads would not map to the ancestor, but instead to the barcoded mutants or neutral strains. See Abreu et al. [27] for full details.

### 4.5 Whole-genome sequencing and ploidy assay

We sequenced 115 mutants from the three-environment evolutions in total; 64 from Glu →H →N, and 51 from Glu →N →H. We used the Illumina DNA Prep kit (ILMN DNA LP (M) Tagmentation (96 samples, IPB)) to prepare samples for whole-genome sequencing, but carried out reactions at 1/5 the volume in the manufacturer’s protocol. We then sequenced these libraries with Admera Health, using 2 *×*150 paired-end sequencing on Illumina NovaSeqX machines.

We followed the same pipeline as in Abreu et al. process FASTQ files: we mapped reads to the S288C genome using bwa, and indexed the BAM file using samtools. SNPs and small indel variants were called using HaplotypeCaller from GATK. Samples were filtered by coverage and alignment and ancestral genotypes and mitochondrial variants genotypes were excluded, classified as homozygous or heterozygous, and annotated using snpEff. Copy number variants were called using samtools and bedtools, selecting for sites with 3x the average coverage or more. Variants appearing in more than 15 samples were excluded as they could be ancestral variants, and results were checked visually using IGV.

We used a benomyl assay [1] to measure the ploidy of the mutants we whole-genome sequenced. We measured the growth of mutants over several days in YPD + 20 µg mL^*−*1^ benomyl, which inhibits the growth of diploid strains substantially more than haploid strains. Plates were imaged and manually classified as haploid or diploid. See Abreu et al. [27] for further details.

### 4.6 Fitness Inference

We adapted the fitness inference method from Ascensao et al. (2023) and Ghosh et al. (2026) [38, 63] for fluctuating environments, and detail the statistical model and inference method below, following similar notation to Ghosh et al. We wish to infer the fitness relative to the ancestor of each strain in each component of a fluctuating cycle, say X →Y→ Z →X We define the fitness of the ancestor strain to be 0, and model the frequency of a mutant i in environment e_*t*_ between time points t and t + 1, x_*i,t*_ as

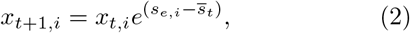

where s_*e,i*_ is the fitness of strain i in e_*t*_, and 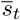 is the mean population fitness at time t. This corresponds to an exponential growth rate difference of s_*e,i*_ between strain i and the ancestor strain if the growth process were simple exponential growth.

We model the noise in read counts from biological sources (stochasticity in growth and due to bottlenecks) and measurement (library preparation and sequencing) as a counting process with a negative binomial distribution, or equivalently each read is sampled from an overdispersed Poisson distribution:

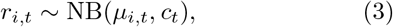

where µ_*i,t*_ is the expected number of reads, and c_*t*_ is the dispersion parameter, such that

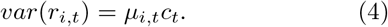

The expected number of reads µ_*i,t*_ is simply the expected frequency of strain at time t multiplied by the total number of reads R_*t*_, so

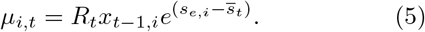

In order to infer s_*e,i*_, we must first infer s_*t*_, the mean fitness of the population, and c_*t*_, the dispersion parameter.

We use the fact that there are 60 “neutral” lineages included in our competition pool, which by definition have fitness 0, in order to measure the mean population fitness and the time specific variance.

For a “neutral” lineage with fitness 0, its frequency at time t, denoted 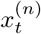 is

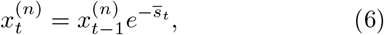

which can be rewritten to give

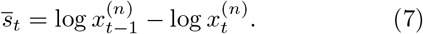

We estimate 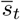 as

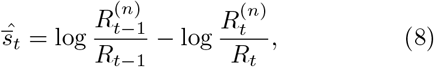

where 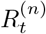 is the total reads summed over all neutral lineages at time t.

We can use the variability in the behavior of the neutral lineages to estimate the dispersion parameter at each time point. Variation between neutral lineages should be due to a combination of genetic drift and technical and measurement noise.

We first consider variation due to genetic drift by considering the Langevin equation for a neutral (fitness 0) strain,

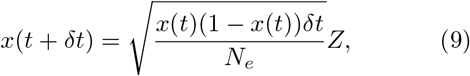

where Z is the standard normal variable, x(t) is the frequency, and N_*e*_ is the effective population size. We apply a variance stabilizing transformation to x(t), and define

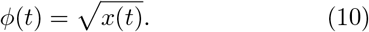

We can now see that this transformation removes the dependence of δϕ = ϕ(t + δt) *−* ϕ(t) on x(t):

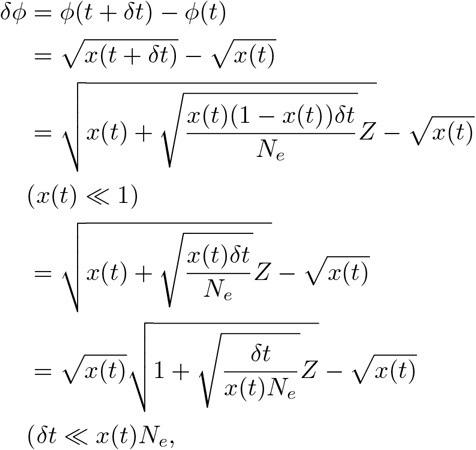

using the Taylor expansion 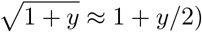

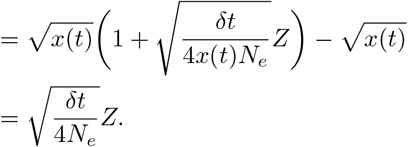

In the above derivation, we assume that δt *≪* x(t)N_*e*_, i.e. the change in time is small compared to the total number of cells in the lineage. Thus, the variance in δϕ due to genetic drift is var 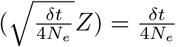. Given that the sources of measurement or technical noise should be independent at each time point, we can simply sum the variance due to these sources when writing an expression for var 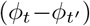. The variance due to drift should accumulate linearly over time. Therefore,

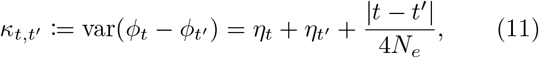

where η_*t*_ is the variance in the reads due to measurement or technical error at timepoint t. Assuming that N_*e*_ does not vary significantly over each passage, we can fit 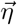 and 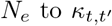 across all time point pairs, which can be estimated as variance across neutral (ancestor) lineages. We only use lineages which had a starting frequency above 10^*−*5^.

We can now relate this back to the dispersion parameter for each time point, to get

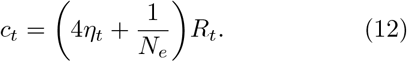

We can now write the likelihood for a particular value of the number of reads assigned to a strain i at time point t, r_*i,t*_, given a particular value of s using the pdf of the negative binomial function:

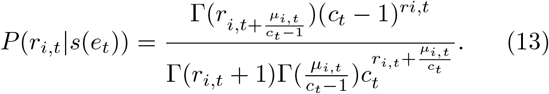

The likelihood of the read counts over the whole time series is then given by P(**r**_**i**_| **s**) = Π_*t*_ P(r_*i,t*_ |s(e_*t*_)). We estimate s jointly with frequency (x) by minimizing the negative log likelihood function over the corresponding time points via gradient descent using the `L-BFGS-B` method from `scipy.optimize.minimize.`

### 4.7 Inference of f_*t*_, f_*t−*1_, and δ from two-environment data

We fit two-environment data to the model s(X, Y → X) = f_*i,t*_(X) + f_*i,t−*1_(Y ), which we are able to do because we have multiple conditions where the same environment is either the previous or the current environment. We use fitness data on 5 environments and all nine pairwise fluctuating environments (excluding data from the Lac/H fluctuating environment due to poor replicate-to-replicate correlations). Each of the fluctuating environments have fitness measurements in two environment components, resulting fitness measurements for each mutant in 23 environments total.

We fit this model independently for each mutant. We use one-hot encoding to represent the environmental conditions for each data point; each fitness measurement has two variables associated with it, the identity of the current and previous environments respectively. For static environments, the previous environment would be the same as the current environment. As the previous and current environment are categorical variables taking one of five values, we one-hot encode as five binary variable. For instance, the fitness of a mutant in Glu preceded by Gal would have 1s associated with it in the “current Glu” and “previous Gal” variables, and 0 with all other variables.

Fitness in environment X preceded by Y, s(X, Y→ X) is modeled as

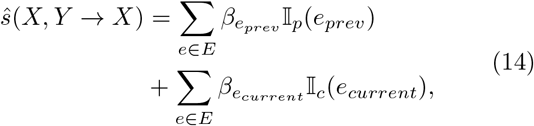

where s(X, Y →X) is the fitness of the strain in environment X preceded by environment Y, E is the set of all environments in the dataset, and the indicator functions I_*p*_ and I_*c*_ encoding environmental conditions are defined as follows:

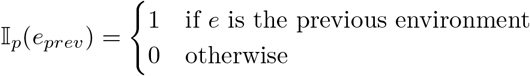

and

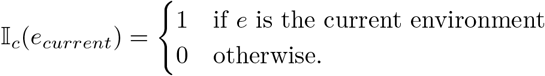

We then fit the fitness value to these binary variables jointly across all environments in the training set, using ridge regression to avoid instability due to multicollinearity.

This model minimizes the loss function

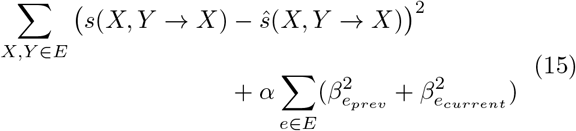

over βs, where α is the regularization parameter,. We use a leave-one-out cross validation to fit a model for each data point separately, training on the fitness data of that mutant on all other environments to generate a predicted fitness value for the held-out environment (Figure 3B).

Because of the binary encoding of environments, the resulting coefficients for each environmental effect are interpretable as the literal fitness contribution of the environment on a mutant’s fitness. The prediction error from this model is jointly due to interactions between the environments and noise in the data.

The regularization coefficient α used in these fits is set to 0.6 by optimizing mean-squared error, generated from LOO-CV over the entire dataset. MSE and our broad results are not very sensitive to α values between 0.1 - 1 (Figure S7A).

Finally, we estimated error in measurement using the standard error of the mean, and subtracted it from the prediction error to estimate δ. If the SEM exceeded prediction error, then the estimate of δ is set to be 0.

As a control, we simulated data with purely additive, partly non-additive and fully non-additive behavior and ran the additive regression model on this synthetic data. We find that this method predicts fitness well in the case of pure additivity, and infers additive coefficients well even under non-additive effects. This method generally overestimates the magnitude of δ, and systematically overestimates correlations between previous and current coefficients by∼ 0.1 and underestimates coefficients by ∼0.1 between two previous or two current coefficients. We correct for this bias in Figure 3H, and show raw correlation values in Figure S7D. See Supplementary Materials for further details.

### 4.8 Estimation of True Positive Two-Cycle Memory

We compare three-environment fitness against the expected fitness from two-environment measurements using Welch’s t-test on the three replicates for each measurement. We find that 11.39% of three-environment measurements have nominal p-values less than 0.05, but none of these survive Benjamini-Hochberg FDR correction correcting across all comparisons (n = 14640) at the threshold α = 0.05.

We estimate the number of true positives by analyzing the empirical distribution of p-values by using Storey’s π_0_. We calculate π_0_ for intermediate p-values at threshold λ, which is defined as

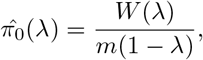

where W (λ) is the number of tests with p-values larger than λ, and m is the total number of tests. 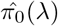 should approach the proportion of true nulls for large p-values, or as λ approaches 1. We use a cubic spline to fit 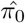 to λ between 0.05 and 0.96, sampled in increments of 0.01 and extrapolate to estimate 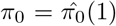, or the fraction of true negatives. We find π_0_ = 0.691 (Figure S10M, blue line), which suggests that 31% of the comparisons are true positives.

## Acknowledgments

The authors thank Alexandra Khristich, Nathan Hill, Mathis Leblanc, Jean Vila, Rachel Porter, Alexander Sastokas, Jan Skotheim, Benjamin Good, and all members of the Petrov and Cremer Labs for helpful comments and discussions. Part of the computing for this project was performed on the Sherlock cluster, which is supported by the Stanford Research Computing Center and Stanford University. This work was supported by the Stanford Graduate Fellowship and Stanford Interdisciplinary Graduate Fellowship (to S.M.), NIH Grant No. R01GM149611 (to J.C.), NIH NIGMS Grant No. 5R35GM118165-07 (to D.A.P.), and NSF-GRFP (to O.M.G.). D.A.P. is a Chan Zuckerberg Biohub - San Francisco Investigator.

