## Supplementary Materials for "Order invariance of fitness makes evolution predictable despite strong history-dependent effects"

### Supplementary Information for “Order invariance of fitness makes evolution predictable despite strong history-dependent effects”

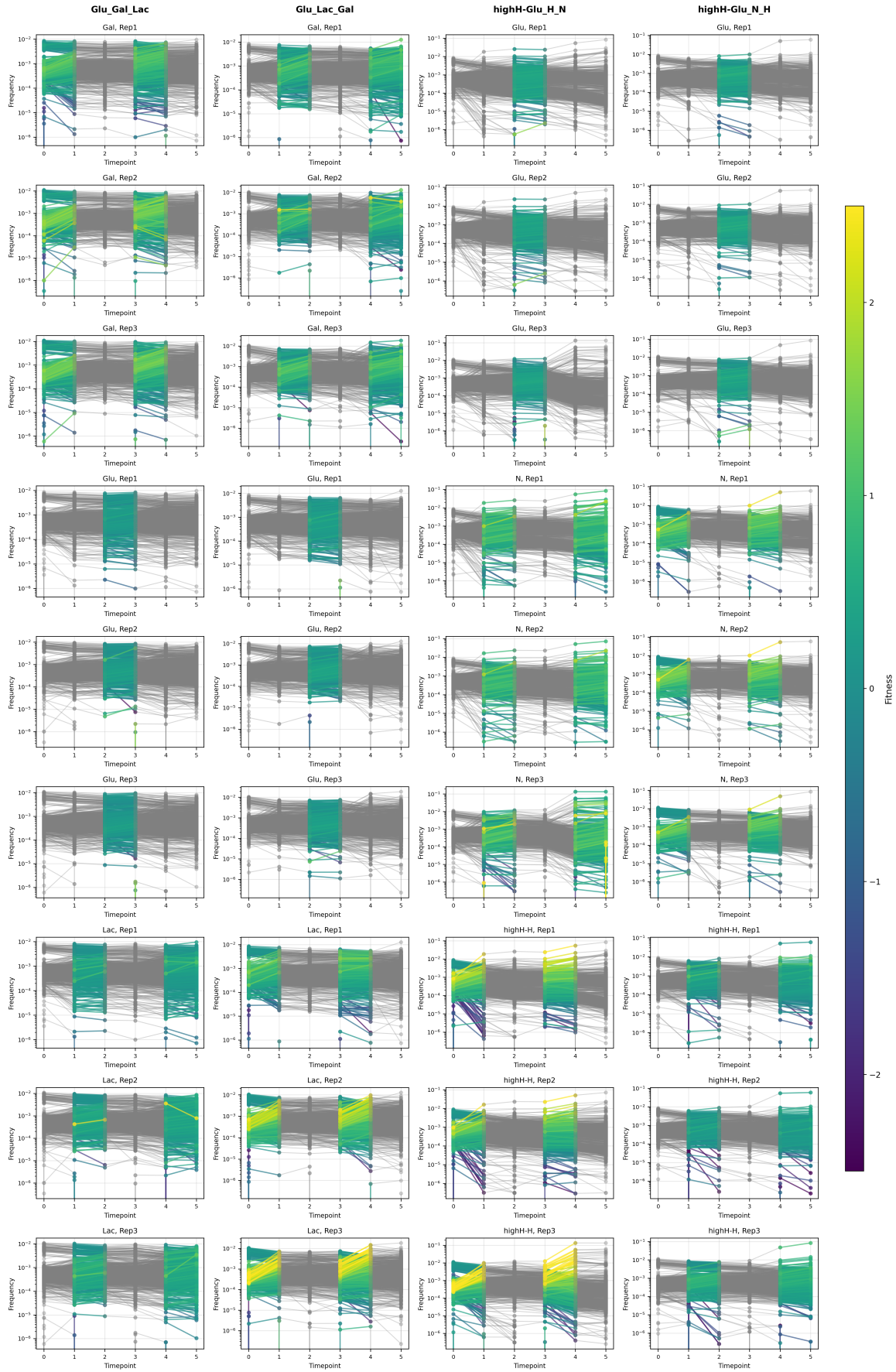

**Figure S1: Frequency trajectories in each environment** Each trajectory represents a single barcode; frequency is simply the number of read-counts assigned to the barcode divided by the total number of reads in the sample. Trajectories are colored by inferred fitness. Plots are grouped by environment (column) and the environment component, with replicates shown next to each other. Lines are colored by the inferred fitness of the barcode in the specific environment component, with the rest of the trajectory shown in gray. The colorbar ranges from -2.5 to 2.5, with fitness values outside this range colored by the extreme values of the colormap; 0.1% of the data lies outside this range.

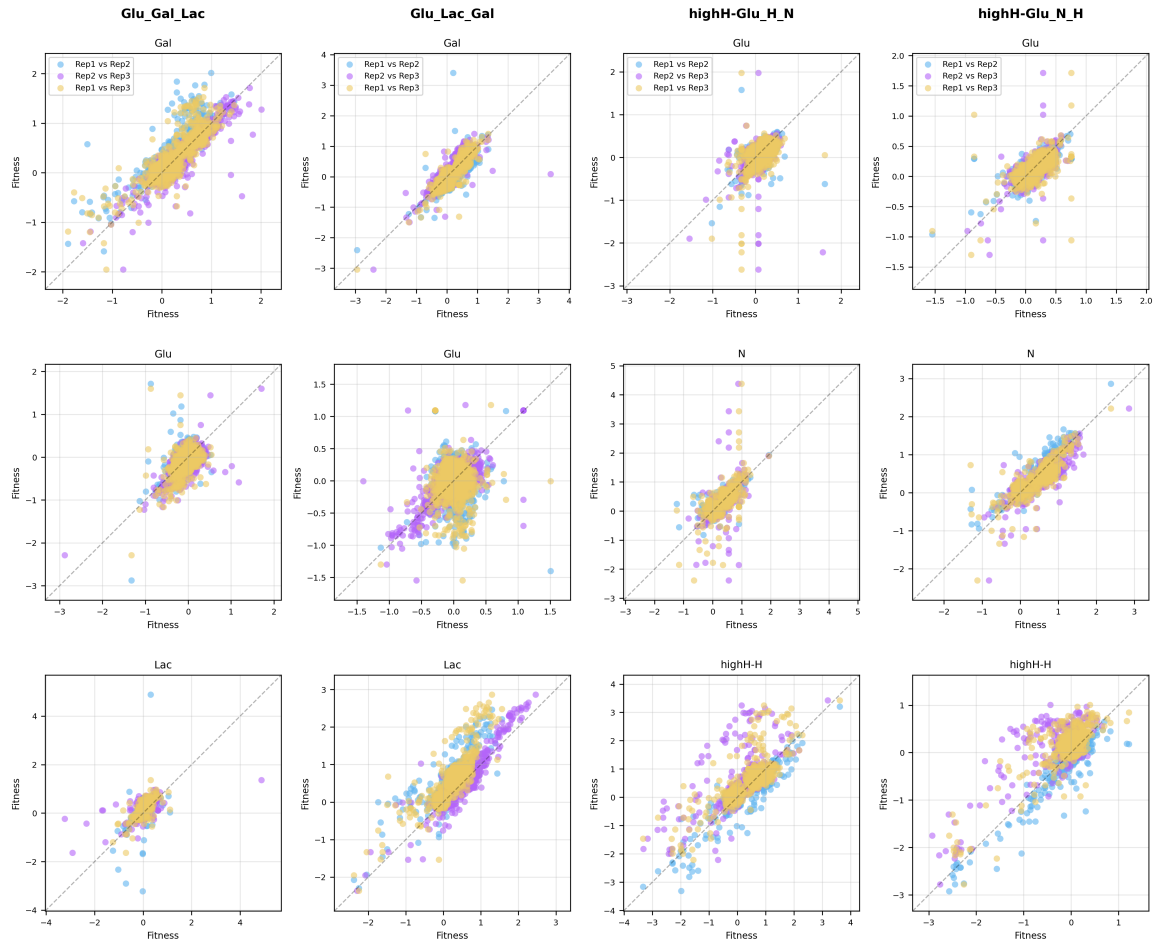

**Figure S2: Replicate-replicate plots for three-environment inferred fitness** Plots showing variation in inferred fitness across each pair of replicates. Points are colored by the replicate-pair labels, and the on-to-one line is shown in grey. Plots are organized by the three-environment cycle (columns).

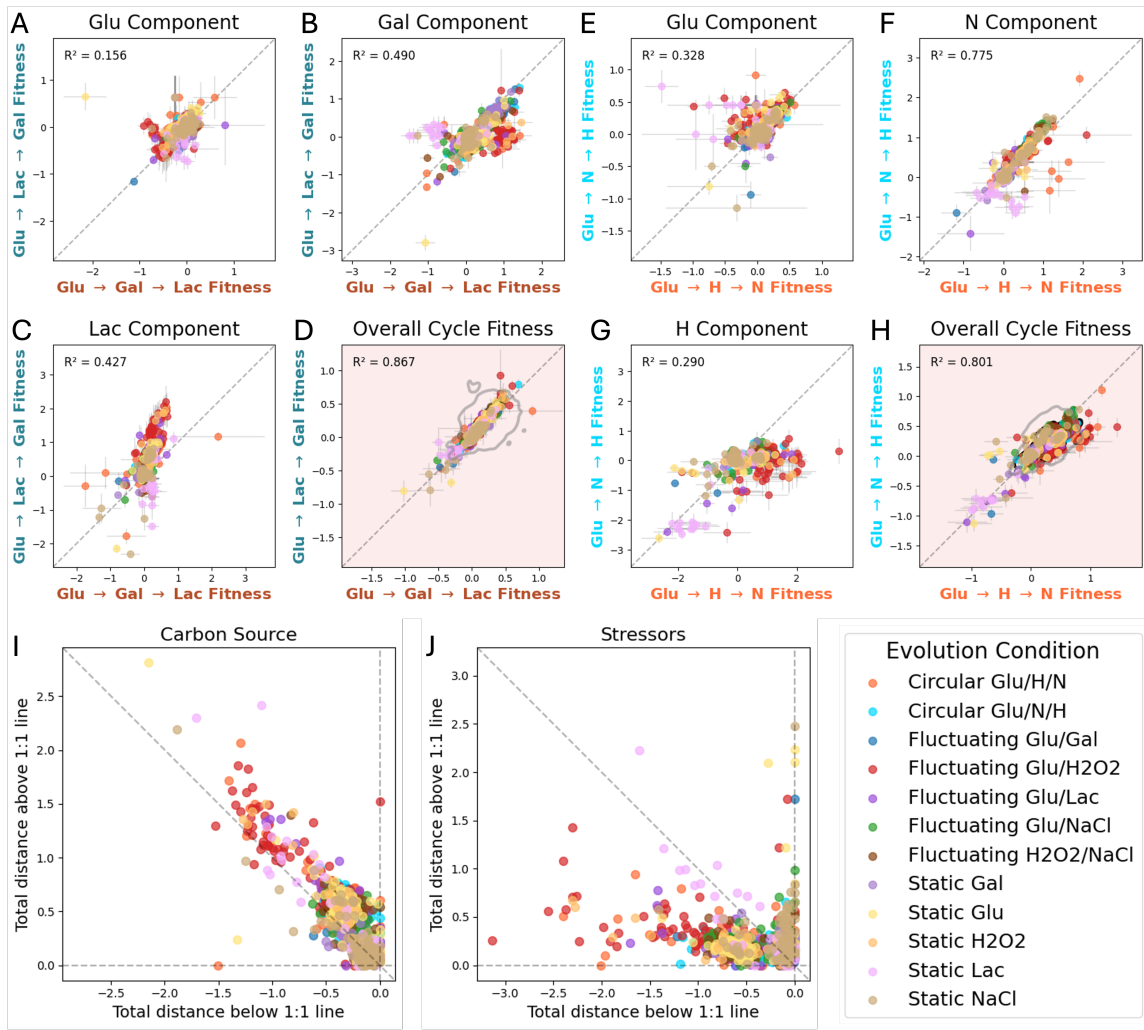

**Figure S3: Three-Environment Fitness Measurements Colored by Source (A-C)** Fitness in each component of the carbon cycle shown in each direction, **(D)** overall fitness in the carbon cycle, **(E-G)** Fitness in each component of the stressor cycle, **(H)** overall fitness in the stressor cycle. A-H correspond to Figure 2 in the main text. **(I-J)** Plots showing the total degree of under-performance in each component of the three-environment cycle, calculated by sum of the Euclidean distances of each barcode below the one-to-one line in each component against over-performance in each component, similarly calculated as the sum of distance above the one-to-one line. The anti-correlation between these values shows that fitness advantages in any one direction of the cycle trades-off with disadvantages in other components.

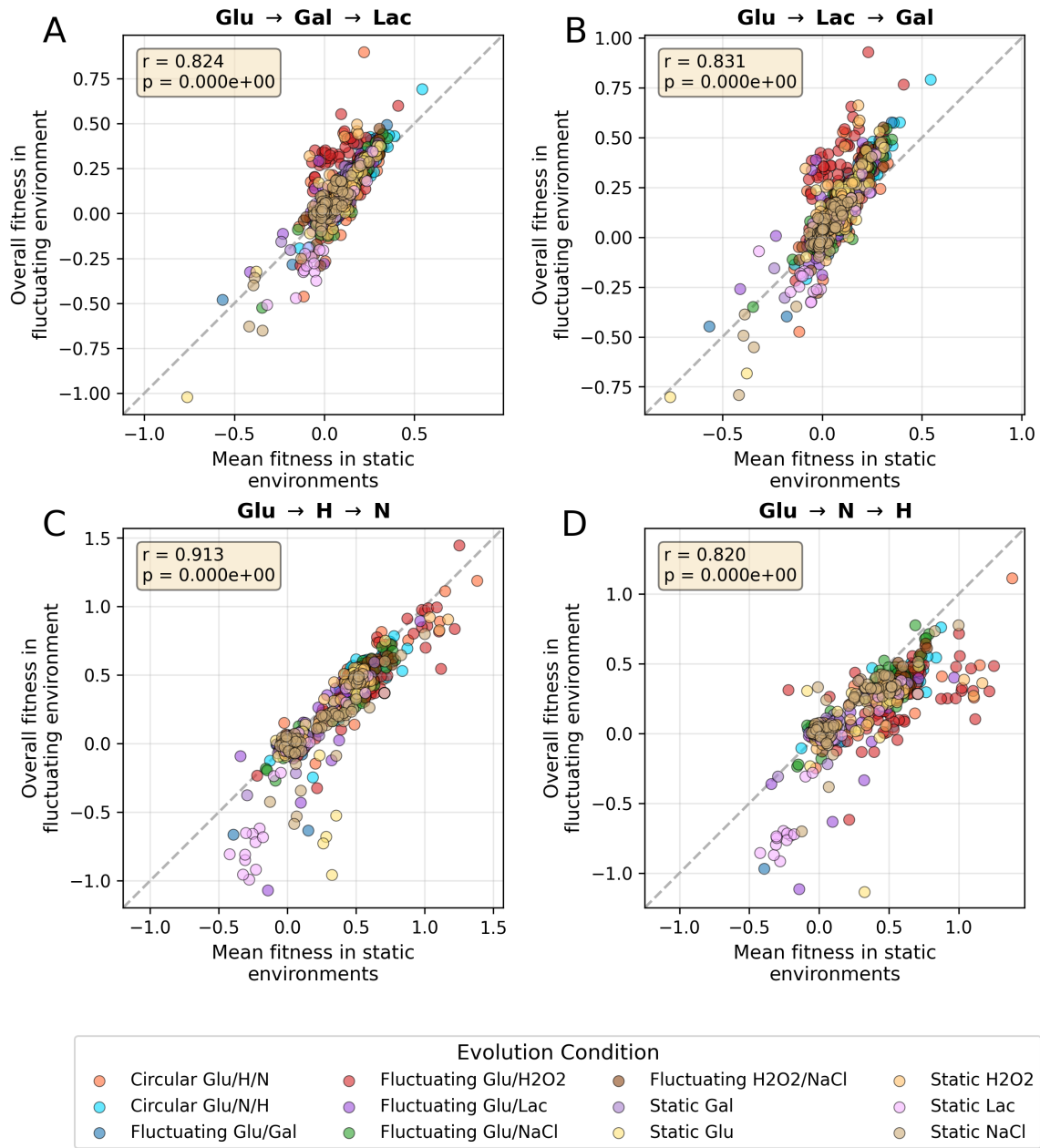

**Figure S4: Additivity of Fitness in Three-Environment Cycles (A-D)** Mean fitness in static environments which comprise the three-environment cycle (Glu, H, N for the stressor cycle and Glu, Gal, Lac for the carbon cycle) is plotted against the overall fitness in the fluctuating environments, which is calculated as the mean fitness inferred in each component of the cycle. Strains are colored by their source environment. $r$ and $p$-values (calculated using a $t$-distribution with `scipy.stats.pearsonr`) are shown with each plot. Mean fitness in each of the static environments is generally a good predictor of overall fitness in the three-environment fluctuating cycle, despite large history-dependent effects.

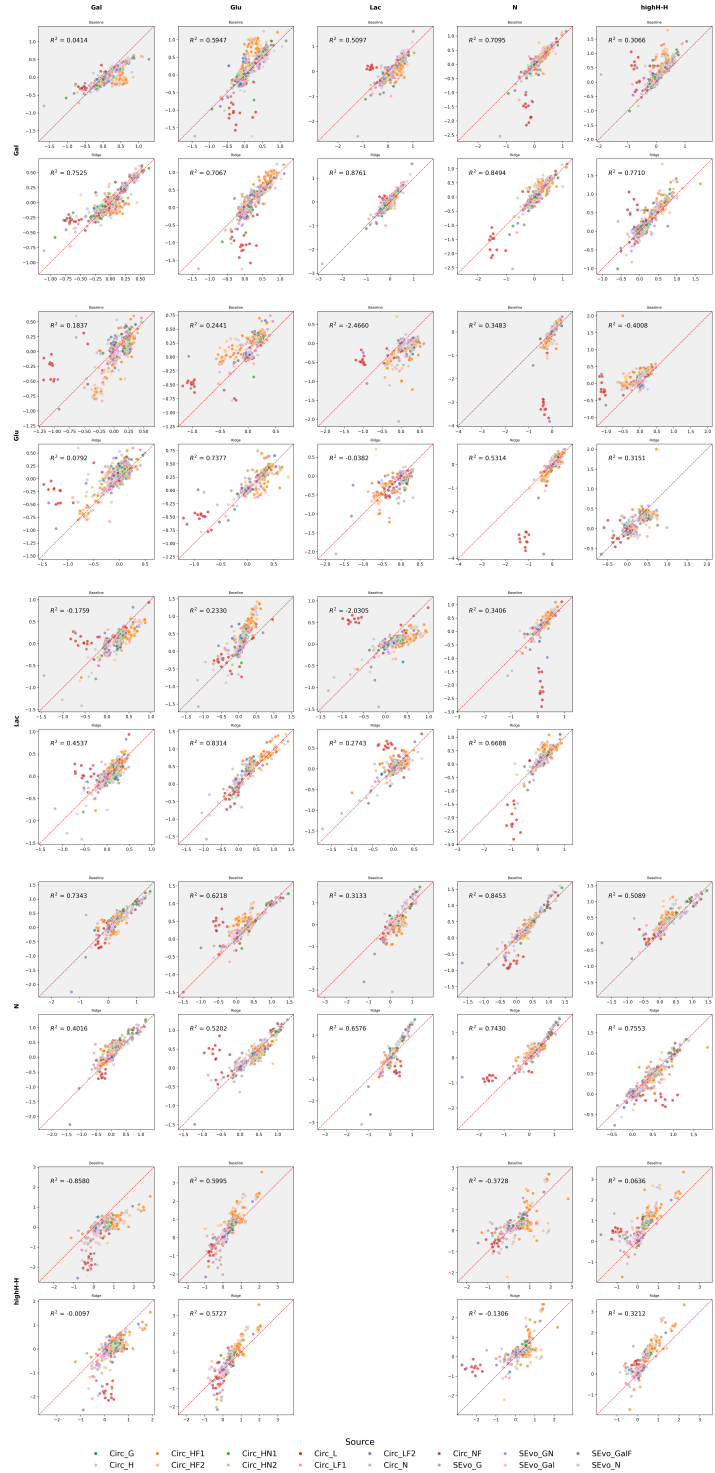

**Figure S5: Out of sample predictions for each environment component** Out of sample predicted fitness from the model fitting $f_t$ and $f_{t-1}$ as described in the main text, compared to measured fitness. Points are colored by source environment, and are organized by the previous environment (row), current environment (column). The baseline prediction taking mean fitness in all environments with the same current environment (assuming there is no environmental memory) are shown in gray, results of the predictions with the additive model are shown in white. The dashed red line is the one-to-one line.

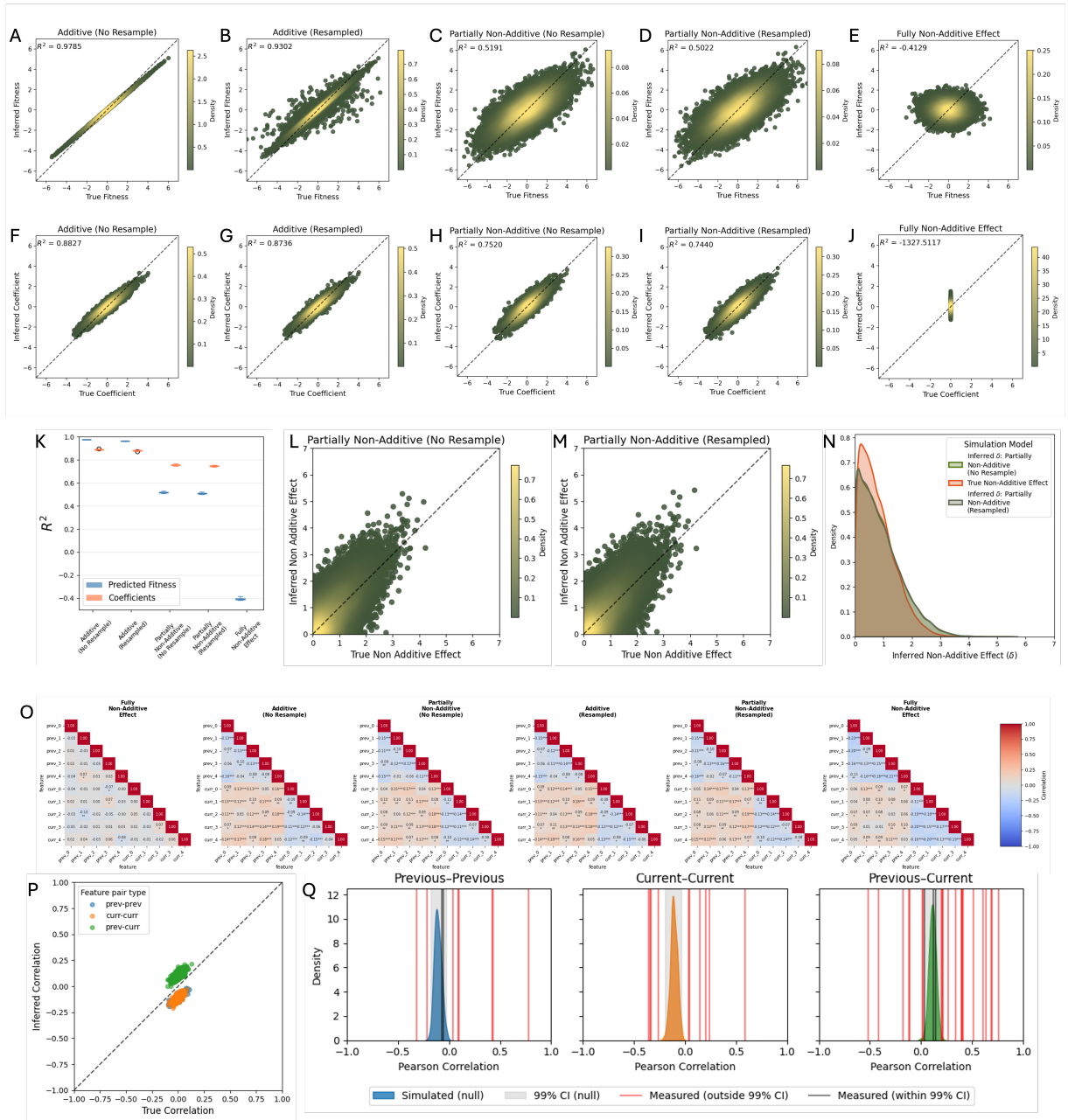

**Figure S6: Simulated model of additivity with one-hot encoding and ridge regression.** (A-E) Predicted fitness compared to true fitness under different models of additivity and noise. (F-J) Inferred additive coefficients ($f_t$ and $f_{t-1}$) compared to true fitness under different models of additivity and noise. (K) $R^2$ values of predicted fitness (blue) and inferred coefficients (orange) compared to truth values. (L-M) Inferred magnitude of $\delta$ values compared to true values with (M) and without (L) added noise. (N) KDE-density plot of true non-additive effect (orange) and inferred $\delta$ with (green) and without (blue) added noise. $\delta$ values are overestimated by this method. (O) True (leftmost) values of correlations between coefficients for one instance of the simulation compared to correlations between inferred coefficients, colored by Pearson correlation values. (P) True correlation values between coefficients compared to inferred coefficient values, showing systematic bias in different types of coefficient pairs. (Q) Distribution of correlations between previous/previous, current/current and previous/current coefficients from 20 runs of the simulation compared to measured correlations. Correlation values outside the 99% confidence interval from the simulated null are shown in red.

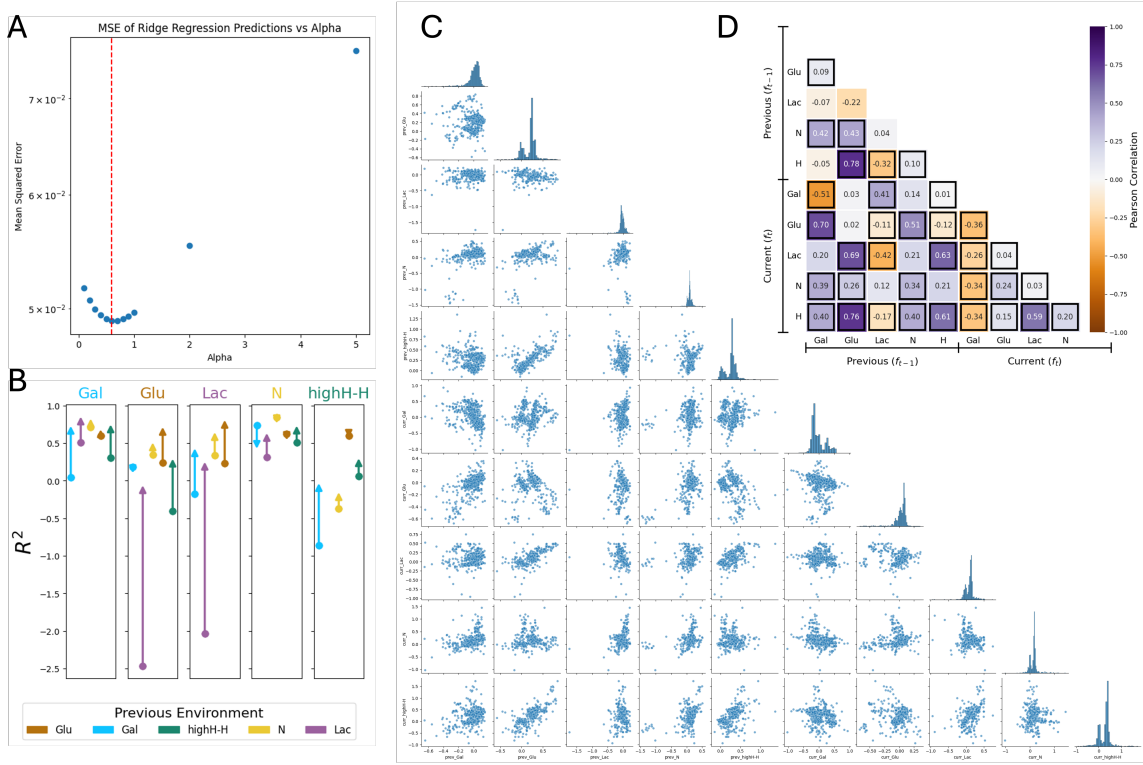

**Figure S7: Empirical results from additive regression model** (A) Mean Squared Error over a range of shrinkage or penalty parameter $\alpha$. MSE is calculated using LOOCV over environments, as with all out-of-sample predictions in this study. Red line at $\alpha = 0.6$ is close to the minimum MSE value, and is used for the data. (B) $R^2$ values of a baseline model, mean fitness in all environments with the same current environment (circles) and $R^2$ from additive regression model (arrow), grouped by the current environment. $R^2$ values increase for most environments, but decrease marginally in one H and two N environments, and decreases substantially in the Gal → N environment. Environments in which the additive model does not improve prediction generally have lower memory, or equivalently, higher baseline $R^2$ values. (C) Pair-plot of previous ($f_{t-1}$) and current ($f_t$) fitness effects. Diagonals show the distribution of each coefficient. (D) Uncorrected correlation values previous ($f_{t-1}$) and current ($f_t$) fitness effects. Correlations that are significant (Permutation test, p-value $\leq 0.05$) and outside the 99% CI of the null distribution from simulations (red in Figure S6P) are highlighted.

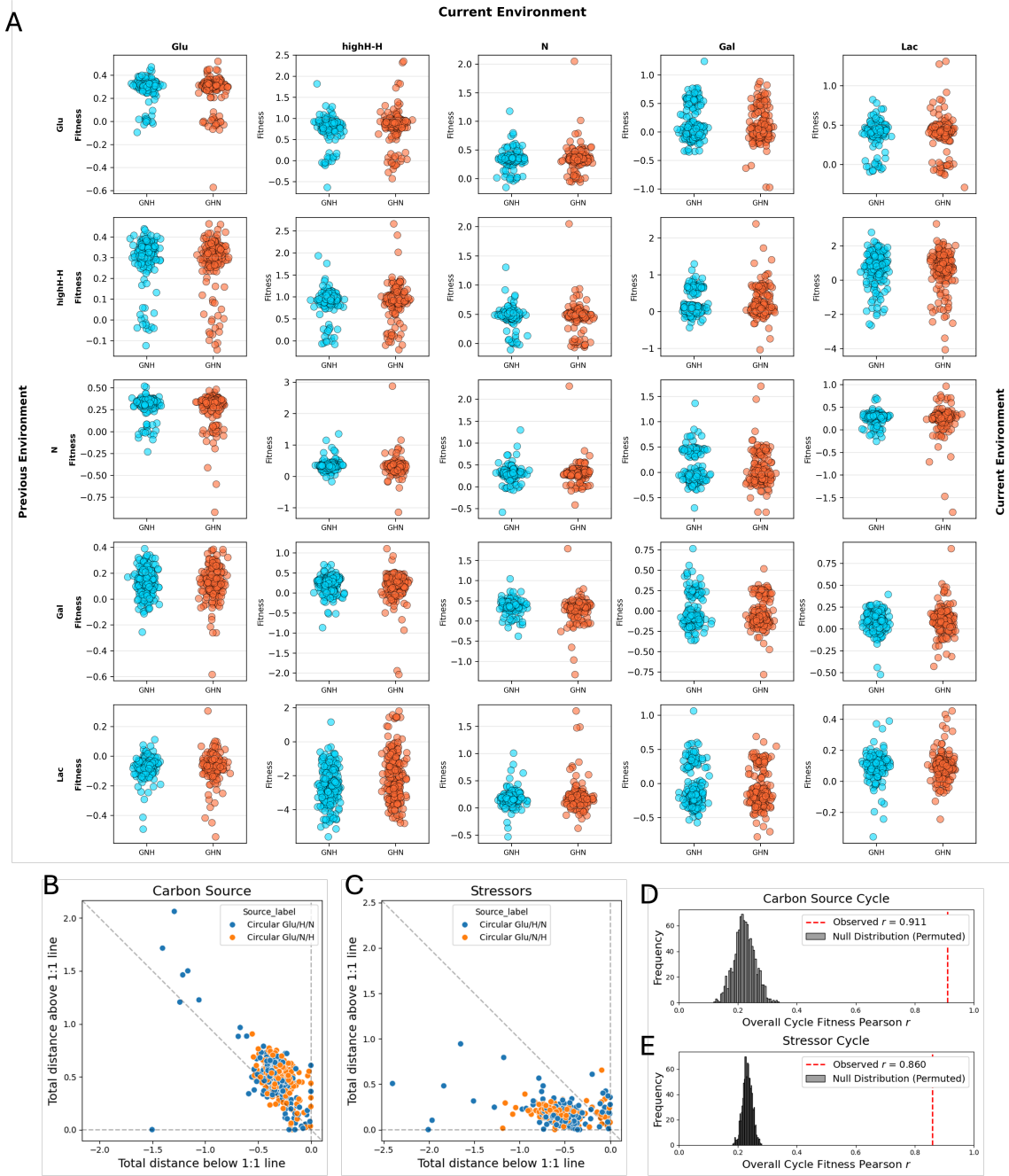

**Figure S8: Fitness of mutants evolved in three-environment cycles in one- and two-environment cycles.** (A) Mutants evolved in Glu → H → N (GHN) are shown in orange, mutants evolved in Glu → N → H (GNH) is shown in blue. Plots are arranged by current environment (columns) and previous environment (rows). The x-axis is jittered for visibility. (B-C) Plots showing the total degree of under-performance in each component of the three-environment cycle, calculated by sum of the Euclidean distances of each barcode below the one-to-one line in each component against over-performance in each component, similarly calculated as the sum of distance above the one-to-one line. The anti-correlation between these values shows that fitness advantages in any one direction of the cycle trades-off with disadvantages in other components. (D-E) Correlation of overall fitness of three-environment mutants. Similar to other mutants, the Pearson correlation of the overall fitness of three-environment mutants in both carbon and stressor cycles far exceeds the null. The correlation values 0.91 and 0.86 are marginally smaller than the the Pearson correlation of overall fitness of other mutants, which is 0.931 and 0.895 in the carbon and stressor cycles respectively.

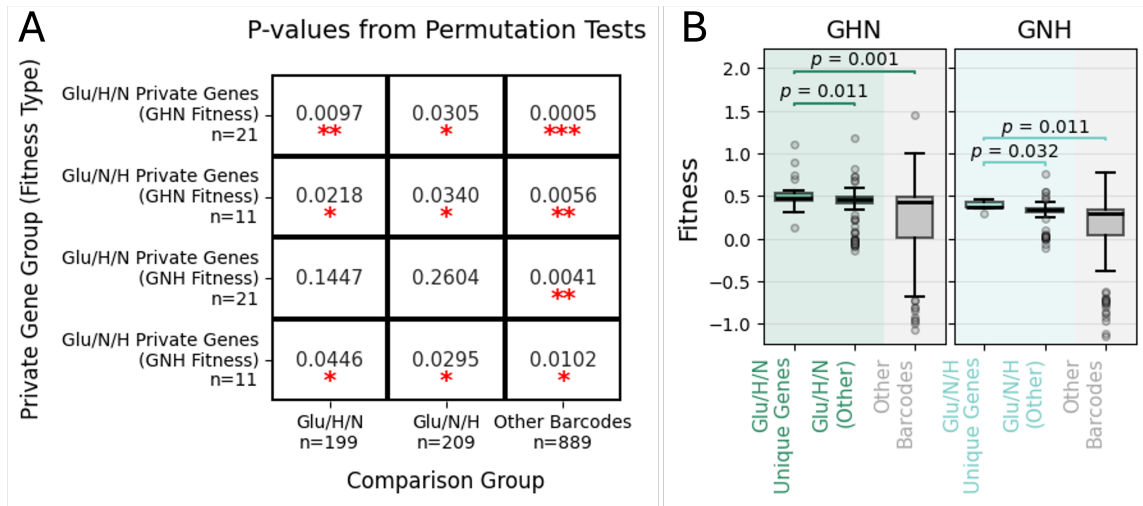

**Figure S9: Strains with multi-hit “private” genes unique to three-environment cycles have higher fitness in three-environment cycles.** “Private genes” are defined as genes hit multiple times in the three-environment cycles which were not hit in any other environments. **(A)** p-values (Permutation test) from comparison of the fitness in GHN or GNH of strains with private genes in the two cycles to other strains from the three-environment cycles or strains from other environments. **(B)** Fitness of strains with private genes have significantly higher fitness compared to other strains from the three-environment cycles and strains from other environments in both the GHN and GNH cycle. P-values are calculated using a permutation test. The highest fitness values of strains with private genes do greatly exceed those of other strains, but strains with private genes nearly always have fitness substantially above zero in the three-environment cycles.

| Category | Gene | Annotation |
| --- | --- | --- |
| GHN private (n = 15) | APM3 | functions in transport of alkaline phosphatase to the vacuole |
|  | DUR3 | Plasma membrane transporter for both urea and polyamines |
|  | ENT5 | involved in clathrin recruitment and traffic between the Golgi and endosomes |
|  | LTP1 | Protein phosphotyrosine phosphatase of unknown cellular role; activated by adenine |
|  | MDM10 | Moonlighting protein component of both the ERMES and SAM complex |
|  | PUF2 | PUF family mRNA-binding protein |
|  | RPO31 | RNA polymerase III largest subunit C160; part of core enzyme |
|  | SEF1 | DNA-binding transcription factor; exerts very weak regulatory role in the expression of TCA cycle genes |
|  | SSC1 | Hsp70 family ATPase; constituent of the import motor component of the Translocase of the Inner Mitochondrial membrane |
|  | SWA2 | Auxilin-like protein involved in vesicular transport |
|  | UBP14 | Ubiquitin-specific protease; specifically disassembles unanchored ubiquitin chains |
|  | VMR1 | Vacuolar membrane protein; involved in multiple drug resistance and metal sensitivity |
|  | LAM5 | Putative sterol transfer protein |
|  | YTA7 | Chromatin segregase involved in DNA replication |
|  | ZRT1 | High-affinity zinc transporter of the plasma membrane; responsible for the majority of zinc uptake |
| GNH private (n = 6) | CYR1 | Adenylate cyclase; required for cAMP production and cAMP-dependent protein kinase signaling |
|  | HIR3 | Subunit of the HIR complex; a nucleosome assembly complex involved in regulation of histone gene transcription |
|  | JJJ2 | Protein of unknown function |
|  | NNK1 | Protein kinase; implicated in proteasome function; interacts with TORC1, Ure2 and Gdh2 |
|  | PAU11 | Protein of unknown function; mRNA expression appears to be regulated by SUT1 and UPC2 |
|  | RTT10 | WD40 domain-containing protein involved in endosomal recycling |
| GHN + GNH private (n = 6) | ERO1 | Thiol oxidase required for oxidative protein folding in the ER; essential for maintaining ER redox balance |
|  | INO80 | Nucleosome spacing factor |
|  | NUP1 | FG-nucleoporin component of central core of the nuclear pore complex |
|  | REC114 | Protein involved in early stages of meiotic recombination |
|  | RPI1 | ATPase of the 19S regulatory particle of the 26S proteasome |
|  | YDR537C | Dubious open reading frame, overlaps with PAD1 |

**Table S1: Private genes by category, with annotation.** GHN private genes are multi-hit genes targeted only in GHN; GNH private genes are multi-hit genes targeted only in GNH; GHN + GNH private genes are multi-hit genes targeted in GHN or GNH, but never in other environments

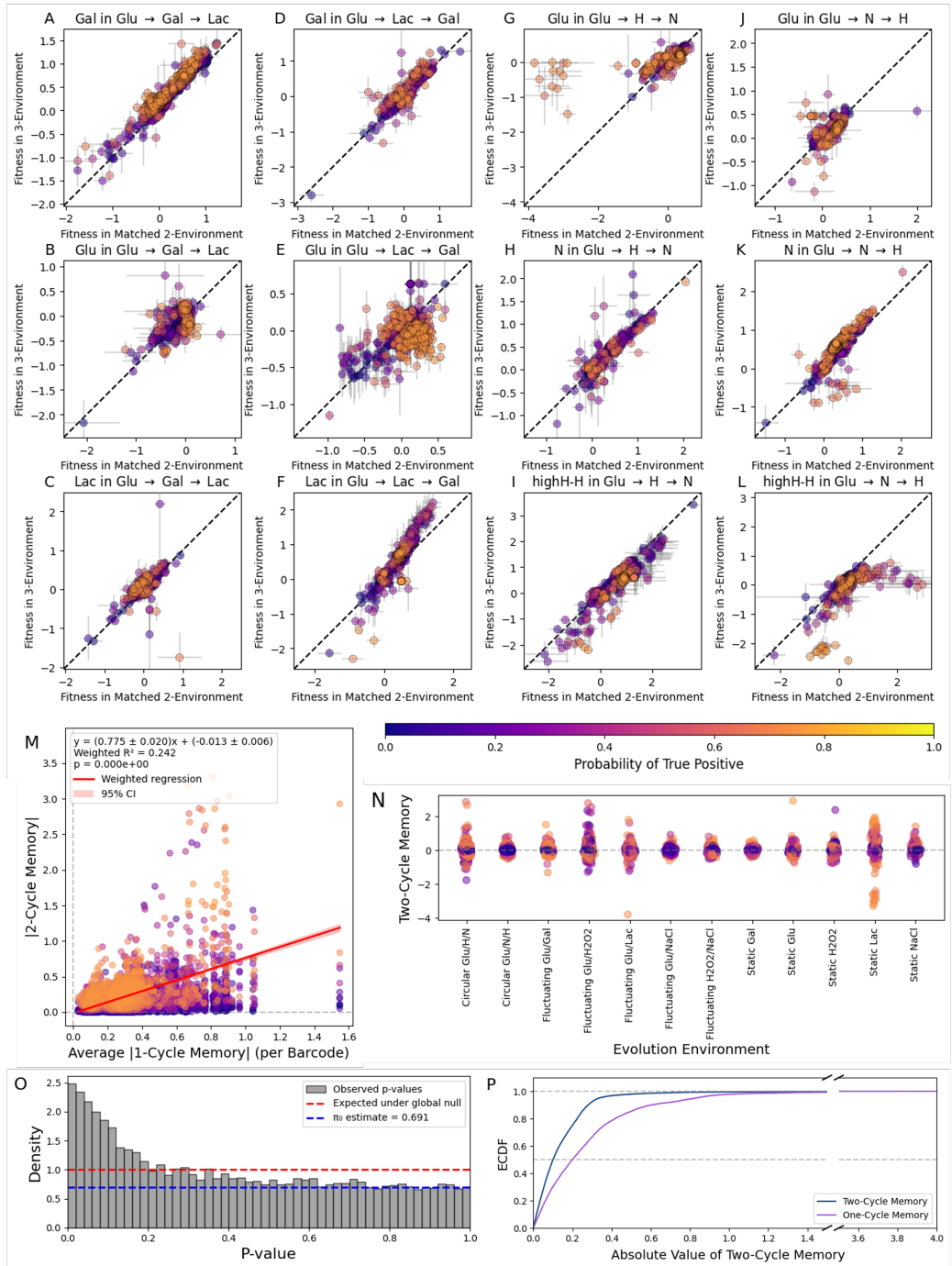

**Figure S10: Environmental memory persists beyond one growth cycle (A-L)** Fitness in each three-environment component is compared to fitness in the two-environment component which has the same preceding (and current) environment. For instance, in **A** fitness in the Gal environment in Glu → Gal → Lac is compared to Gal fitness in the Glu/Gal environment. These conditions share current (Gal) and previous (Glu) environments, but differ in the environment two cycles before the measurement. Each point represents the mean fitness of a mutant in the given environment, error bars are given by the standard error of the mean (SEM). Points are colored by the likelihood of true positive difference in fitness from two-environment estimate (using Welch's t-test and Storey's $\pi_0$; see text for further details). Plots are labeled by environment component and cycle. **(M)** Average one-cycle memory across environments plotted against magnitude of two-cycle memory. We see a weak but significant correlation ($R^2 = 0.24$, $p < 10^{-15}$). **(N)** Magnitude of two-cycle memory by evolution environment of the mutant. A few mutants from static lactate, Glu/H, and Glu → H → N show particularly large two-cycle memory effects. **(O)** Histogram of p-values (Welch's t-test), showing enrichment for low p-values compared to the null expectation (red). Storey's $\pi_0$ (blue) indicates the expected density of the true nulls, or the noise floor. Density above this line can be interpreted as signal. Points in **A-L** are colored by the likelihood of being above this line in their bin (using 100 bins). **(P)** ECDF plot showing the magnitude of memory effects from one cycle (purple) and two cycles (blue). The x-axis is clipped from 1.5 to 3.5 (diagonal lines). One-cycle memory appears to be generally greater in magnitude than longer-term two-cycle memory.

#### 1 Fitness in three-environment cycles is structured by evolution condition

Examining the fitness measurements in the three-environment cycles further, we find that patterns in the three-environment fitness data is structured by the evolutionary history. To illustrate this we colored strains in Figure S3 by their evolution condition.

For instance, for the stressor cycle strains that have evolved in static H<sub>2</sub>O<sub>2</sub> (orange) or fluctuating Glu/H<sub>2</sub>O<sub>2</sub> (red) are fitter in H when preceded by Glu (Glu → H → N direction) than when preceded by N (Glu → N → H direction) (Figure S3G). This suggests that they have evolved specifically to the Glu to H transition. Interestingly, these mutants have asymmetric fitness in the carbon cycle, where their higher fitness in the galactose environment of these mutants in the Glu → Gal → Lac direction is off-set by their lower fitness in this direction in lactate. On the other hand, mutants evolved in static galactose (purple, Figure S3A-H) do not show a large difference in fitness in the two directions in any of the three-environment components, and in fact cluster around a fitness of 0 in all of the environments in these cycles with the exception of galactose.

These patterns are also consistent with cross-protection effects in stressor conditions. In the Glu → H → N direction, hydrogen peroxide is the first component, and has the largest absolute fitness values, whereas in the Glu → N → H direction, N has the largest spread in fitness values. This is most readily visible by comparing the spreads in the axes in the hydrogen peroxide conditions in the stressor cycle (Figure S3G). Performance in the first stressor having a disproportionately large effect is consistent with the model that cells activate a general stress response when in any stressor (high or H<sub>2</sub>O<sub>2</sub>), thus leaving all strains on more equal footing due to cross-protection going into the next environment. In this framework, fitness in these fluctuating environments depends on the degree to which cells can grow in a stressful environment coming from a less stressful environment, which is consistent with the model that fitness in these environments comes primarily from lag phase.

#### 2 Null model controls for statistical estimation of additivity

As a control, we simulated data under different degrees of additivity and levels of noise, and ran the same regression model, where fitness $s(X, Y \rightarrow X) = f_t(X) + f_{t-1}(Y) + \delta(Y \rightarrow X)$ to check the extent to which $f_t$, $f_{t-1}$, and $\delta$ values can be correctly inferred by this method.

We simulate the fitness in 5 component environments and all 23 possible combinations with 900 mutants, as in the dataset shown in Figure 3 over 20 iterations. In order to do this, we simulate additive fitness components for each mutant $i$ and environment component. For pairs of environments components, we simulate additional non-additive terms. We simulate noise by drawing SEM values from the empirical distribution of SEM values with replacement from the data. We then draw three replicate measurements from a normal distribution centered around the simulated fitness with standard deviation $\sigma$ as $\sqrt{n}SEM$, where $n = 3$ is the number of replicates. We draw values for each variable as follows:

$$\begin{aligned} f_{i,t}(X) &\sim N(0, 1), \\ f_{i,t-1}(Y) &\sim N(0, 1), \\ \delta(Y \rightarrow X) &\sim N(0, 1), \\ s_{add,i}(X, X \rightarrow Y) &= f_{i,t-1}(X) + f_{i,t}(Y), \\ s_{nonadd,i}(X, X \rightarrow Y) &= f_{i,t-1}(X) + f_{i,t}(Y) + \delta(Y \rightarrow X), \\ SEM_i(X, X \rightarrow Y) &\sim \text{Uniform}(S), \\ s_{add,i,j}(X, X \rightarrow Y) &\sim N(s_{add,i}(X, X \rightarrow Y), 3SEM_i(X, X \rightarrow Y)), \\ s_{nonadd,i,j}(X, X \rightarrow Y) &\sim N(s_{nonadd,i}(X, X \rightarrow Y), 3SEM_i(X, X \rightarrow Y)), \\ \bar{s}_{add,i}(X, X \rightarrow Y) &= \sum_j \bar{s}_{add,i,j}(X, X \rightarrow Y), \\ \bar{s}_{nonadd,i}(X, X \rightarrow Y) &= \sum_j \bar{s}_{nonadd,i,j}(X, X \rightarrow Y), \end{aligned}$$

where $s_{add,i}(X, X \rightarrow Y)$ and $s_{nonadd,i}(X, X \rightarrow Y)$ are additive and non-additive fitness, $s_{add,i,j}(X, X \rightarrow Y)$ and $s_{nonadd,i,j}(X, X \rightarrow Y)$ are replicate $j$ of the sampled additive and non-additive fitness, and $\bar{s}_{add,i}(X, X \rightarrow Y)$ and $\bar{s}_{nonadd,i}(X, X \rightarrow Y)$ are sample means.

We then run the regression on each of five versions of the data and compare inferred coefficients and fitness against the true values, which is shown across all 20 instances of the simulation in Figure S6A-K. In general, inferred fitness is very well correlated with true fitness under the additive model (Figure S6A, $R^2 = 0.979$), but is systematically smaller in magnitude, which is expected when using a ridge regression method which penalizes the magnitude of the coefficients. Inferred coefficients are slightly less tightly correlated with their true values (Figure S6F, $R^2 = 0.883$); this occurs because the same predicted fitness, which is simply the sum of two coefficients can be achieved by many combinations of coefficients, and by penalizing large coefficients, ridge regression pushes coefficients to be smaller in magnitude than their true values.

When adding noise in the form of sampling noise (Figure S6B,G), non-additive effects (Figure S6C,H), or both (Figure S6D,I) the model retains some degree of predictive power for fitness, and a large degree of predictive power for additive coefficients. The model has a sharp drop-off in prediction of fitness from the additive model to adding large non-additive effects, as might be expected, but has a much more modest drop-off in predicting additive coefficients in the partially additive model (Figure S6K). The model with only non-additive terms (Figure S6 E,J) fails to predict inferred fitness, as expected. Together, these simulations show that this regression method can infer additive coefficients well even under a large degree of non-additive effects, with a small bias towards inferring smaller coefficients than their truth values.

Next, we compare inferred delta values to true non-additive effects (Figures S6L-N). Estimates $\delta$ is less tightly correlated with true values than the predicted fitness and coefficients, and are systematically slightly over-estimated (Figure S6N), suggesting that the claim that $\delta$ is small from the data is conservative.

Next, we check the extent to which spurious correlations between coefficients can be generated by the regression method. Figure S6O (leftmost) shows the true correlation values between the coefficients, and other plots show the correlation between inferred coefficients for one replicate of the simulation, and Figure S6P compared true coefficient for this same data under the partially non-additive resampled model, which is closest to the empirical data. The regression method appears to systematically increase the correlation of previous-current pairs by $\sim 0.1$ and decrease the correlation of previous-previous or current-current pairs by $\sim 0.1$. Intuitively, this is because the previous and current environments are paired together, and previous-previous or current-current environments are never paired together.

Using all 20 replicates of the simulation, we construct a null distribution of the Pearson correlations under this model. This distribution captures the bias and spread due to this inference method, as well as spurious correlations from sampling a finite number of mutants where the underlying Gaussian distributions are uncorrelated. We compare correlation values to this null, and consider values outside the 99% confidence interval of this distribution to be significant (red lines in S6Q). We correct for bias in Figure 3H simply by subtracting the mean value of the Pearson correlation of the null distribution from the corresponding type of coefficient pair.

##### 3 Additivity in three-environment cycles can occur if and only if effects of each environment on future fitness are additive also

We have shown empirically that there is a high degree of “additivity” in fitness in fluctuating cycles; for many strains overall fitness in the three-environment cycle is similar across both directions (Figure 2) and overall fitness is well predicted by the mean fitness in static environments in both two and three-environment cycles ([Abreu et al., 2024], Figure S4). However, fitness in individual components differs drastically depending on the identity of the previous environment (Figure 2). We can show that additivity only occurs if and only if the effects of an environment on fitness in the current and subsequent time-point can be split additively.

Let’s define the environmental “memory” $m(Y \rightarrow X)$ as the fitness difference in a fluctuating environment, $X$ preceded by $Y$, relative to fitness in the static environment $s(X, X \rightarrow X)$, or $s_X$:

$$m(Y \rightarrow X) = s(X, Y \rightarrow X) - s(X, X \rightarrow X). \quad (1)$$

From additivity in two-environment cycles, we have that

$$s_X + s_Y = s(X, Y \rightarrow X) + s(Y, X \rightarrow Y) \quad (2)$$

$$= s_X + s_Y + m(Y \rightarrow X) + m(X \rightarrow Y), \quad (3)$$

which is easily rewritten to show that $m(X \rightarrow Y) = -m(Y \rightarrow X)$, which is shown directly in [Abreu et al., 2024]. Now, if we consider a three-environment cycle, overall fitness $\bar{s}$ in the $X \rightarrow Y \rightarrow Z \rightarrow X$ cycle is $\frac{1}{3}[s(X, Z \rightarrow X) + s(Y, X \rightarrow Y) + s(Z, Y \rightarrow Z)]$. Now, using the additivity in a three-environment cycle, we can see:

$$\bar{s} = \frac{1}{3}[s(X, Z \rightarrow X) + s(Y, X \rightarrow Y) + s(Z, Y \rightarrow Z)] = \frac{1}{3}[s_X + s_Y + s_Z] \quad (4)$$

$$\implies m(Z \rightarrow X) + m(X \rightarrow Y) + m(Y \rightarrow Z) = 0 \quad (5)$$

$$\implies m(X \rightarrow Y) + m(Y \rightarrow Z) = -m(Z \rightarrow X) \quad (6)$$

$$\implies m(X \rightarrow Y) + m(Y \rightarrow Z) = m(X \rightarrow Z) \quad (7)$$

$$(8)$$

For a function $f$, if $f(X, Y) + f(Y, Z) = f(X, Z)$ then there exist functions $f_1$ and $f_2$ such that $f(X, Y) = f_1(X) + f_2(Y)$. In order to see this, Let’s consider the function $g(A) = f(A, x_0)$, where $x_0$ is an arbitrary value. If $f(X, Y) + f(Y, x_0) = f(X, x_0)$, then this can be rewritten as $f(X, Y) = g(X) - g(Y)$. Therefore, memory $m(Y \rightarrow X)$ can be expressed as $f_1(X) + f_2(Y)$. Fitness in a fluctuating environment, $s(X, Y \rightarrow X)$, can therefore be written as $s_X + m(Y \rightarrow X) = s_X + f_1(X) + f_2(Y)$, or in our notation, $f_t(X) + f_{t-1}(Y)$.

The reverse direction is more straightforward to see - if fitness can be decomposed into the effect of the previous and current environments, $s(X, Y \rightarrow X) = f_t(X) + f_{t-1}(Y)$, then overall fitness in a cycle of environments $X_1, X_2 \dots X_n$ can be written as $\sum_i f_{t-1}(X_i) + f_t(X_i) = \sum_i s_{X_i}$.
